# Mechanical stress initiates and sustains the morphogenesis of wavy leaf epidermal cells

**DOI:** 10.1101/563403

**Authors:** Amir J Bidhendi, Bara Altartouri, Frédérick P. Gosselin, Anja Geitmann

**Affiliations:** Institut de Recherche en Biologie Végétale, Département de sciences biologiques Université de Montréal, Montreal, Quebec, H1X 2B2, Canada; Department of Plant Science, McGill University, Macdonald Campus, 21111 Lakeshore, Ste-Anne-de-Bellevue, Québec H9X 3V9, Canada; Laboratoire de Mécanique Multi-échelles, Département de génie mécanique, École Polytechnique de Montréal, C.P. 6079, Succ. Centre-ville, Montreal, Quebec, H3C 3A7, Canada

**Keywords:** Morphogenesis, plant cell mechanics, feedback loop, stress and strain, buckling, leaf epidermis, pavement cell, pectin, cellulose, cell wall, finite element analysis

## Abstract

Plant cell morphogenesis is governed by the mechanical properties of the cell wall and the resulting cell shape is intimately related to the respective specific function. Pavement cells covering the surface of plant leaves form wavy interlocking patterns in many plants. We use computational mechanics to simulate the morphogenetic process based on experimentally assessed cell shapes, growth dynamics, and cell wall chemistry. The simulations and experimental evidence suggest a multistep process underlying the morphogenesis of pavement cells during tissue differentiation. The mechanical shaping process relies on spatially confined, feedback-augmented stiffening of the cell wall in the periclinal walls, an effect that correlates with experimentally observed deposition patterns of cellulose and de-esterified pectin. We provide evidence for mechanical buckling of the pavement cell walls that can robustly initiate patterns *de novo* and may precede chemical and geometrical anisotropy.

**Highlights:**

- A multistep mechano-chemical morphogenetic process underlies the wavy pattern of epidermal pavement cells.
- Microtubule polarization is preceded by an event that breaks mechanical isotropy in the cell wall.
- Mechanical models simulate the formation of wavy cell shapes, predict buckling of the cell walls and spatially confined variations in the mechanical properties of leaf epidermal cells.
- Stress/strain stiffening following the buckling of the cell walls constitutes a crucial element in a positive feedback loop forming interlocking pavement cells.
- Polarization of cortical microtubules, cellulose microfibrils, and de-esterified pectin occur at the necks of wavy pavement cells, matching the *in silico* prediction of cell wall stiffening.

## Introduction

Differentiation of plant cells in the apical or lateral meristems begins with simple spherical or polyhedral bodies produced by cell division. The cells formed at the shoot and root apical meristems are approximately cubic whereas those deriving from lateral meristems tend to be longitudinal and brick shaped—all possess a relatively simple geometry. During differentiation of plant organs, tissue-dependent cell types emerge that exhibit a kaleidoscopic array of different morphologies depending on location and function (Mathur, 2004). In plant and fungal cells, the morphodynamics of differentiating cells is intimately linked to the mechanics of the extracellular matrix—the cell wall. While shaping processes are regulated by the cytoskeleton, they are not mechanically driven by it, since the forces generated by cytoskeletal arrays are too small to act against the wall (Money, 2007). The plant cell wall regulates the shape of the plant cell by balancing the intrinsic turgor pressure and external mechanical constraints through its compliance. The turgor pressure propelling cell expansion is a scalar acting uniformly on all cellular surfaces. Therefore, to grow into complex shapes, plant cells must meticulously modulate the local mechanical properties of the wall at subcellular scale. This regulatory mechanism is important since, unlike animal cells, morphogenetic processes in plant cells are typically irreversible.

The wall of growing plant cells, the primary wall, is a composite material comprising several types of polysaccharides, proteins, ions and water. Two main structural components of the primary cell wall are pectins and cellulose microfibrils (Bidhendi and Geitmann, 2016; Cosgrove, 2015, 2018). Cellulose microfibrils are generally recognized as the main load-bearing component of the cell wall that confer anisotropy (Anderson et al., 2010; Baskin, 2005; Burgert and Fratzl, 2009; Crowell et al., 2011), whereas pectin chemistry is mostly recognized in the context of the local stiffness of the wall matrix (Bidhendi and Geitmann, 2016; Braybrook and Peaucelle, 2013; Carter et al., 2017; Giannoutsou et al., 2016; Torode et al., 2017). Both cellulose microfibrils and pectins are thought to direct the local shaping of cells, but hitherto these concepts have mostly been investigated in cells with simple shapes such as pollen tubes (Fayant et al., 2010), root hairs (Shaw et al., 2000), trichome branches (Yanagisawa et al., 2015), or cells of root and shoot epidermis (Baskin, 2005; Bou Daher et al., 2018; Peaucelle et al., 2015). How plant cells form complex shapes is poorly understood (Geitmann and Ortega 2009).

Some of the most intriguing manifestations of plant cell morphogenesis occur at the surface of leaves. Formation of interlocking protrusions and indents in leaf epidermal pavement cells is a common trait in many plant species (Vőfély et al., 2019). The resulting pattern of wavy cell shapes resembles a jigsaw puzzle (Figs. 1A). The complex growth pattern of leaf pavement cells makes them an ideal model to study mechanisms underlying the formation of complex shapes in plant cells (Bidhendi and Geitmann, 2018; Jacques et al., 2014; Majda et al., 2017; Mathur, 2004; Sapala et al., 2018; Szymanski, 2014). Various biomechanical concepts have been proposed to explain the formation of lobes in pavement cells (Jacques et al., 2014; Korn, 1976; Korn and Spalding, 1973; Majda et al., 2017; Sapala et al., 2018; Watson, 1942). Hypotheses range from bending of the cell walls resulting from the growth of cells in a confined space, inhibition of pavement cell expansion due to forces from cuticle or inner mesophyll layers to localized outgrowth of the anticlinal cell walls (Korn, 1976). The ‘localized outgrowth’ hypothesis is the most widely accepted explanation for shape generation in pavement cells. It states that regions of localized outgrowth penetrate the neighboring cells (Korn, 1976). However, it does not elucidate how the localized outgrowth is initiated and how its subcellular position is determined. The orchestrated formation of alternate necks and lobes in adjacent cells implies that a mechanism must exist that operates across the walls connecting neighboring cells. In pavement cells, local growth has been correlated with well-organized microtubule bundles in neck regions and accumulation of actin microfilaments at sites of lobes (see Fig. 1B for definition of lobe and neck) both of which had been proposed to be regulated by auxin-mediated antagonistic pathways (Fu et al., 2005; Fu et al., 2002; Xu et al., 2010). New evidence suggests, however, that the cytoskeletal pathways warrant further investigation (Belteton et al., 2017). While details about their specific roles in shaping of pavement cells are elusive, actin microfilaments and microtubules appear to be instrumental in this process (Mathur, 2004, 2006; Smith, 2003; Smith and Oppenheimer, 2005; Zhang et al., 2011). This has been deduced from the pronounced pavement cell shape defects upon pharmacological and mutation-mediated interference with cytoskeletal functioning (Baskin et al., 2004; Baskin et al., 1994; Mathur, 2004). Actin microfilament patches are suggested to promote local outgrowth in lobes through exocytotic delivery of new wall-building materials and wall-loosening agents such as expansins (Cosgrove, 2005; Fu et al., 2002; Panteris and Galatis, 2005; Smith, 2003). Microtubules are generally thought to regulate plant cell wall mechanics by determining the orientation of newly deposited cellulose microfibrils (Crowell et al., 2009; Gu et al., 2010). This is accomplished by guiding the location of insertion and trajectory of cellulose synthase (CESA) enzymes at the surface of the plasma membrane (Hamant and Traas, 2010). Cellulose microfibrils are therefore thought to mimic the orientation of microtubules. Therefore, microtubule orientation is sometimes used as a proxy to infer the localization of cellulose microfibrils (Eng and Sampathkumar, 2018; Fu et al., 2005; Zhang et al., 2011). In an influential paper, Panteris and Galatis (2005) proposed that actin filaments are present mostly in lobe regions promoting expansion, and importantly, that a cortical microtubule array is associated with the anticlinal wall (wall perpendicular to leaf surface, see Fig. 1B) of a neck flaring out under the periclinal wall (Fig. 1C and D). Panteris and Galatis postulate this microtubule array to be mirrored by a local enrichment in cellulose on the neck side of the undulations resulting in stiffening of the cell wall. Mechanical validation of this concept is lacking, however (Jacques et al., 2014). The role of the wall polysaccharides in mediating the mechanical conditions has not been investigated in detail. Further, cellulose orientation has not been directly visualized in pavement cells using fluorescence techniques. In a recent study by Sampathkumar et al. (2014), the effect of pavement cell shape on the stress state of the cell wall and microtubule reorganization was modeled mechanically. However, while this relationship may constitute one step of a patterning mechanism governing development, it does not explain the initiation of polarized shapes and lobe formation *de novo* (Sampathkumar et al., 2014). Importantly, in this study, a convincing causal link is made from cell shape to spatial orientation of microtubules, but the reverse question of the effect of microtubule orientation (or of cellulose microfibrils) on shape development remains unexplored. Recently, Majda et al. (2017) proposed that heterogeneous stiffness distribution in the anticlinal walls underlies border waviness in pavement cells under conditions that cause the wall to be stretched, an original mechanical morphogenetic model that we have assessed in detail (Bidhendi and Geitmann, 2019). Briefly, we showed that this model, which focuses exclusively on the anticlinal cell walls, results in mechanical outcomes that are incompatible with known concepts characterizing pavement cell morphogenesis.

**Fig. 1.**
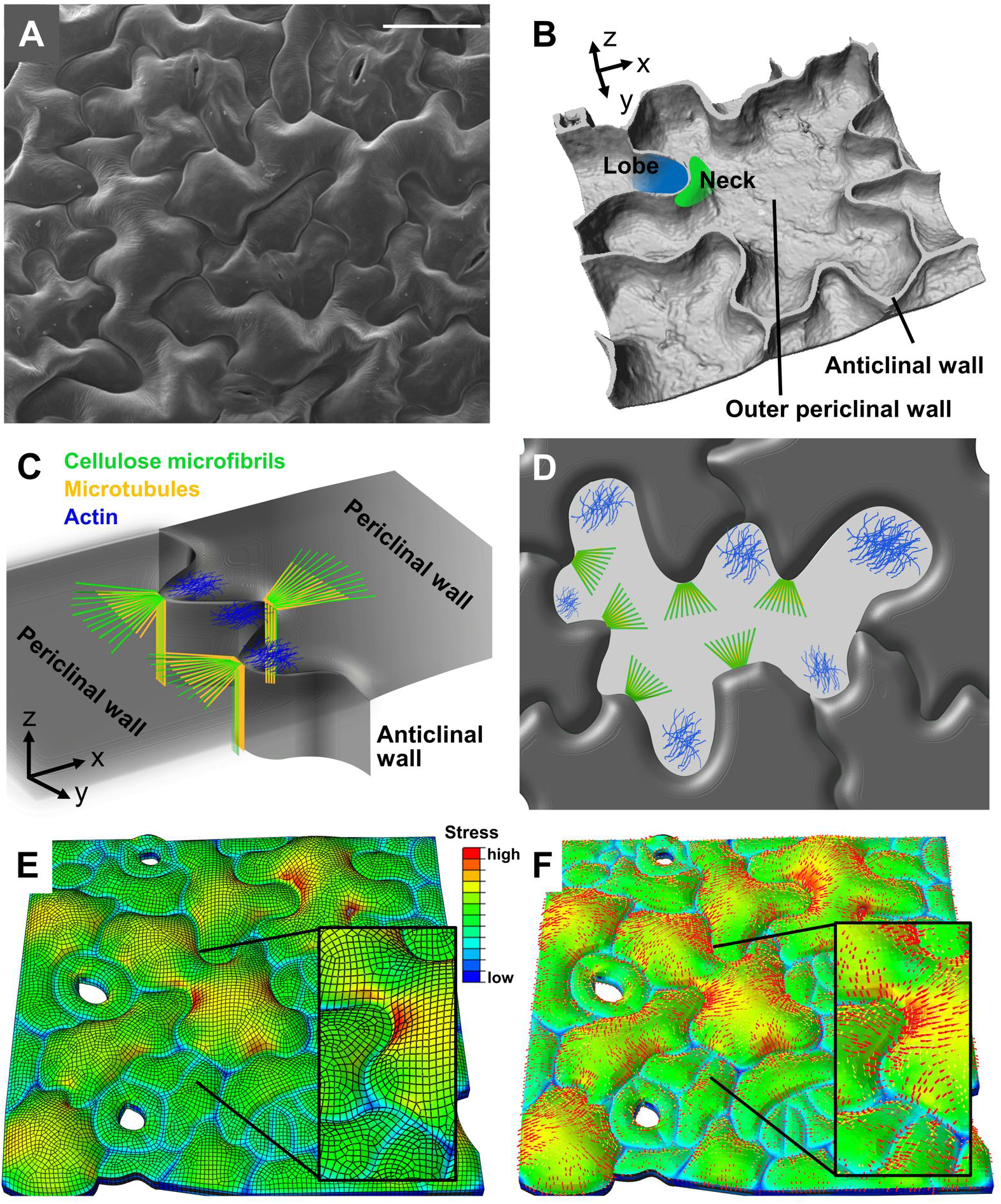
Pavement cells in the leaf epidermis of *Arabidopsis thaliana* form wavy shapes that correlate with epidermis stress pattern. **A)** Scanning electron micrograph showing jigsaw puzzle shaped pavement cells (Scale bar = 30 *μ*m). **B)** 3D reconstruction of confocal microscope z-stack of a pavement cell consisting of the outer periclinal and anticlinal cell walls. C) Local deposition of cellulose microfibrils (red) guided by microtubules (green) on anticlinal and periclinal walls proposed by Panteris and Galatis (2005). Tips of lobes display actin arrays (blue) as shown by (Armour et al., 2015; Fu et al., 2005) **D)** Paradermal view of pavement cells showing localization of microtubules, cellulose microfibrils, and actin microfilaments. **E)** and **F)** Finite element model of a section of epidermis of an *Arabidopsis* cotyledon constructed from the 2D pattern of cell borders extracted from a confocal micrograph. Extrusion of borders was performed to add the anticlinal cell walls. Pressure applied under periclinal walls results in stress in cell walls. **E)** Neck sides of undulations showed higher stresses. At identical pressure, larger cells showed more pronounced out-of-plane bulges than smaller cells. **F)** Orientation of stress field in the epidermis as a result of cell shape and turgidity. Stress fields in periclinal walls radiate in a fan-shaped manner from necks.

Here, we developed 2D and 3D finite element (FE)-based mechanical models to investigate how pavement cell mechanics is involved in forming undulating patterns starting from simple polygonal geometries. We provide experimental validation for the *in silico* predictions by characterizing the spatial distribution of putative mechanical agents including pectin, cellulose microfibrils, and microtubules.

## Results

### Verifiable predictions for cell wall deformation through finite element modeling

Finite element modeling was used to simulate the deformation of the cell wall under the effect of internal pressure (turgor) as the deforming force. The finite element method is a mathematical tool widely used in structural mechanics and allows for solving problems involving complex materials and geometries. The results of the mechanical model were used to interpret the experimentally acquired data on cell wall composition and microtubule arrangement to identify the mechano-structural requirements for the formation of interdigitations in pavement cells. Details of the parameters used in the models are explained in the Supplemental Information. The border undulations in epidermal pavement cells as seen from above correspond to the edge forming the junction between the anticlinal and periclinal walls. We refer to these undulating edges as the *superficial cell borders* (*superficial outer* or *inner* corresponding to outer and inner periclinal walls of a cell, respectively). Since the undulations in the superficial borders of leaf pavement cells correspond to bends in the anticlinal wall, the putative differential expansion of this wall structure has been the primary focus of various experimental or modeling studies (for instance refer to Jacques et al., 2014; Majda et al., 2017; Sapala et al., 2018). We have previously shown that a mechanism based on anticlinal wall stretching proposed by Majda et al. (2017) does not produce undulations when the 3D structure of the cell is considered (Bidhendi and Geitmann, 2019). To further corroborate that exclusive focus on the anticlinal wall is misplaced, we used FE modeling to simulate alternative mechanisms such as differential expansion of the anticlinal wall and differential turgor in neighboring cells (see Notes 1 and 2, Supplemental Information, and results below), but neither approach yielded results relevant to the biological system. Therefore, we moved toward more complete 3D descriptions of pavement cell geometry by considering both anticlinal and periclinal walls.

As a first step, we used the realistic cell geometries of a cotyledon epidermis to confirm that in our hands the modeling approach on a multicellular tissue yields results consistent with those published previously for fully lobed cells. We extracted the cell borders from confocal micrographs of epidermal layers of the *Arabidopsis* cotyledon stained with propidium iodide. 3D cell geometries were created by adding the anticlinal walls at cell borders and pressurizing the cells (Figs. 1E and F). The finite element simulations show that localized stress hotspots develop at the neck sides of undulations on the periclinal walls, consistent with results of Sampathkumar et al. (2014). The simulations also confirmed that, for the same pressure, larger cells bulge out more than smaller cells, as shown by Sapala et al. (2018). High tensile stresses occur at the neck regions of the undulations and in the central regions of periclinal walls. The magnitude of stress in the central regions appeared to correlate with the radius of the largest embeddable circle in that region, as proposed by Sapala et al. (2018). The consistency with these earlier studies provides us with validation of our modeling approach.

### Stress in anticlinal cell walls is highly anisotropic and can become compressive

The presence of the hotspots near cell borders indicates that the bulging of the periclinal wall under turgor may exert a tensile force on the anticlinal walls. To investigate this possibility without the confounding complexity of the lobed cells, two simplified FE models were created representing hexagonally shaped cells with straight borders. One consists of two connected cells that each have 5 cell borders that are free to move in the plane because of lack of neighboring cells (Fig. S1G). The other has a four-cell structure that allows avoiding free edge effects at the central anticlinal wall that is embedded between all four cells (Fig. 2A). Boundary conditions are applied to the corners of the lower periclinal walls to simulate attachment to the underlying mesophyll layer. Upon pressurization of the cells, the periclinal walls bulge out and pull the anticlinal walls inward, reducing the inplane projected cell surface (see Figs. 2D, S1K for the four-cell model and S1G-J for the two-cell model). From the central anticlinal wall in the four-cell model it is apparent that the vertical component of the stress is considerably larger than the in-plane component (*σ_z_* against *σ_x_*, Figs. 2B and C). In other words, finite element modeling indicates that for cell aspect ratios typical for *Arabidopsis* epidermal cells, the stress caused by turgor in the anticlinal wall is highly anisotropic and the mean stress direction is perpendicular to the plane of leaf. Remarkably, the results indicate that under application of turgor pressure, not only is the anticlinal wall exposed to predominant tensile stress in z direction, but the transverse component of the stress (*σ_x_*) can become locally negative (compressive, Fig. 2C). This is due to pressure acting on periclinal walls pulling them out of the plane which in turn results in compression of the anticlinal walls. As compressive forces have the potential to cause buckling, these findings raise the intriguing perspective that, simply based on cell geometry and turgor load, conditions may arise that make the anticlinal walls of pavement cells prone to buckling.

**Fig. 2.**
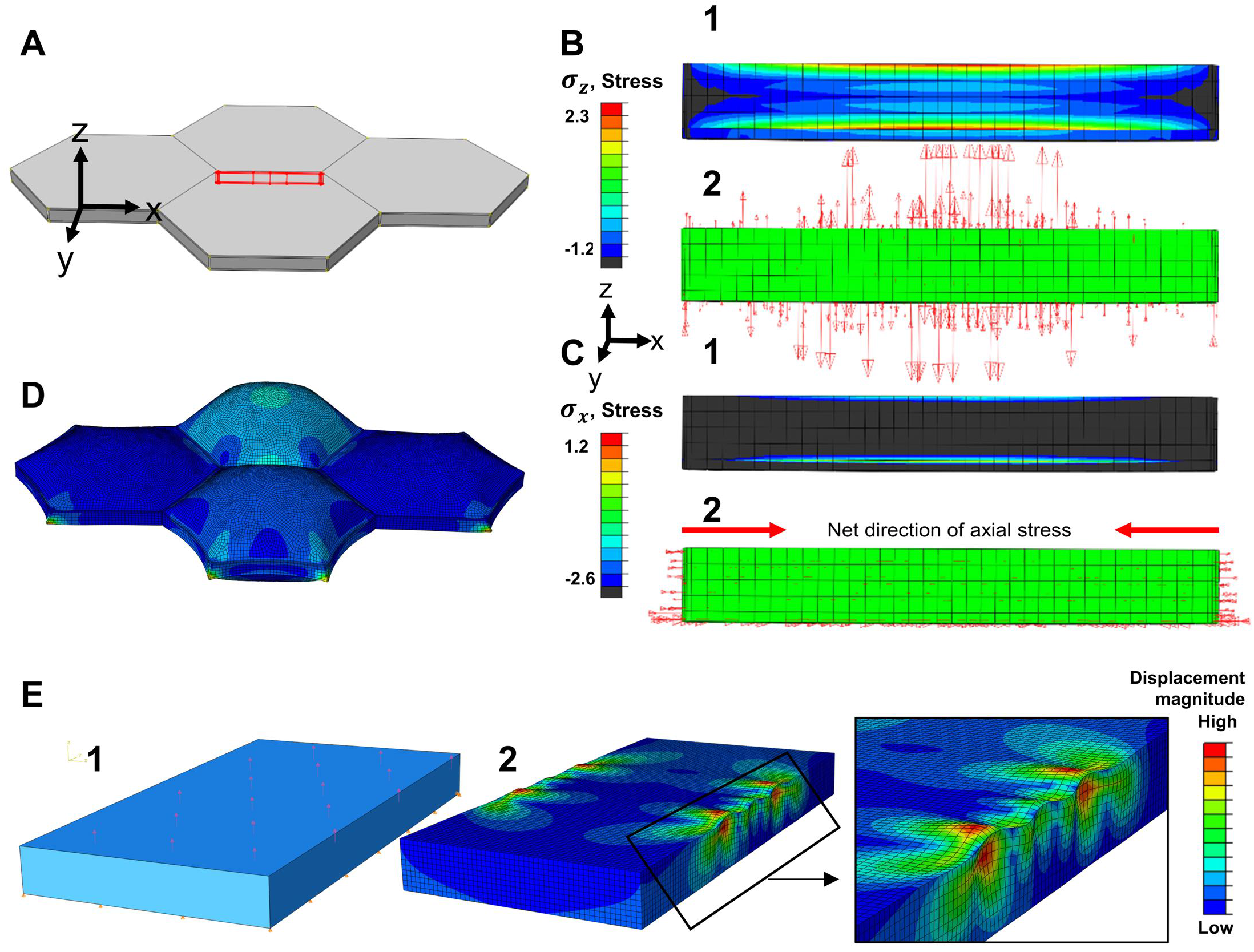
Finite element simulation of stresses and resulting deformations upon load application through turgor pressure. **A)** Structure of four-cell finite element model used to assess stress pattern in the anticlinal wall surrounded by hexagonal cells. The red rectangle marks the anticlinal wall that is shown isolated in B and C to reveal stresses. In this wall, stress components in vertical direction **(B)** are significantly larger than in horizontal direction **(C). B-1)** Color-coded distribution of vertical (z direction) component of wall stress. **B-2)** Arrows indicate the magnitude and direction of the vertical stress. **C-1)** Color-coded distribution of axial (X direction) component of wall stress. **C-2)** Arrows indicating axial stress in horizontal direction revealing that compressive stresses result from the vertical expansion of the cells. Stress values are relative as normalized input values are used in models (see Notes 3 and 4, Supplemental Information). **D)** Model shown in A after turgor application, displaying finite element mesh. Isotropic material properties and equal stiffness in all cell walls are used. Relative turgor values are specified in Fig. S1K. The periclinal cell wall of the cell with the higher turgor pressure forms a pronounced bulge in z direction pulling the anticlinal walls inwards (also see Fig. S1G-K). **E) E-1)** Closed-box model of a brick shaped cell with turgor pressure applied to inner face of the outer periclinal wall. Turgor pressure was not applied to lateral walls as equal pressures were shown in previous models to cancel each other out in a multicell context and only result in compression of the wall thickness. Inner periclinal wall was prevented from out or inward displacement to simulate attachment to mesophyll layer. **E-2)** Buckling mode from the linear buckling analysis demonstrating undulation of the cell border in both periclinal and anticlinal walls. Interestingly, the model indicates that at the location of indentation the periclinal walls bulges out of the plane which is consistent with microscopic observations of pavement cells.

To confirm this possibility, we tested it on an idealized body. For proof of concept, we developed an FE model of a closed-box rectangular cell geometry of similar aspect ratio as pavement cells, subject to internal pressure (Fig. 2E-1; see also Supplemental Information, Note 5). Linear buckling analysis yielded positive eigenvalues. Existence of positive eigenvalues indicates that the structure can buckle under the prescribed loading direction (internal pressure). A buckling mode for this model is depicted in Fig. 2E-2. Interestingly, the model suggests that the periclinal wall at the neck side of indentations bulges out locally (inset in Fig. 2E-2). This is consistent with microscopic observations showing the periclinal walls at the neck sides of undulations bulging out of plane in much steeper manner than at the corresponding lobe side (Fig. 1A and orthogonal view in Fig. 5D (bottom)). These results indicate that the cell walls of pavement cells, with dimensions typical for the *Arabidopsis* cotyledon epidermis (see Fig. S5G and Note 2, Supplemental Information), have the potential to buckle due to cell geometry and internal pressure. It should be noted that in this analysis, we did not incorporate the putative additional compressive forces that may arise due to axial stresses generated in the cell walls as a result of growth-rate mismatch between neighboring pavement cells. These may represent a second origin of buckling behavior.

### Differential stiffness in the periclinal wall can generate cell border undulations

If buckling is indeed an initiating mechanism, it would not be able to create more than rather small deformations. However, these could be an initiating trigger if a subsequent mechanism is able to augment these. Based on a hypothesis postulated by Panteris and Galatis (2005), we sought to assess whether the mechanical properties of periclinal walls play a role in growth and shaping of epidermal cells. We hypothesize that, to form pronounced undulating borders, the mechanical properties of the periclinal wall must vary in alternatingly arranged, spatially confined regions (similar to the organization of microtubules, actin and cellulose illustrated in Figs. 1C, D). We first considered the generation of an isolated bend in the cell border by focusing on a transverse segment including both anticlinal and periclinal walls. In this one-dimensional approach, the periclinal walls of two adjacent cells and a shared anticlinal wall are considered as a T section of beam elements (Fig. 3A). The periclinal walls of the two adjacent cells are assigned different initial stiffness values, and then turgor pressure is applied to the beams (Fig. S2A). The boundary conditions imposed on this model, as with the 2D shell model of Fig. 3B, enable horizontal (in x direction) movement for the end-point of the anticlinal wall (A2) which represents the midway point of the entire anticlinal wall depth. The periclinal end-points (P1 and P2) can only move in the y direction. The point corresponding to the connection of the anticlinal wall with the periclinal walls (A1) may move in all directions. These boundary conditions are applied to all similar models henceforth. The simulations show that upon application of equal pressure in both cells, the periclinal walls bulge out (in the z direction), but the wall with the higher stiffness deforms less than the periclinal wall of the neighboring cell. Consequently, the anticlinal wall moves from the mid-point towards the cell with the stiffer periclinal wall. This displacement of the mid-point connecting the two cells could be considered as the evolution of a protrusion with the lobe being formed by the cell with the lower periclinal wall stiffness (Fig. S2B). The 2D thin shell model of the same situation (Fig. 3B) yields the same results confirming the robustness of the predictions independently of the element type (Figs. 3C, and S2C, D).

**Fig. 3.**
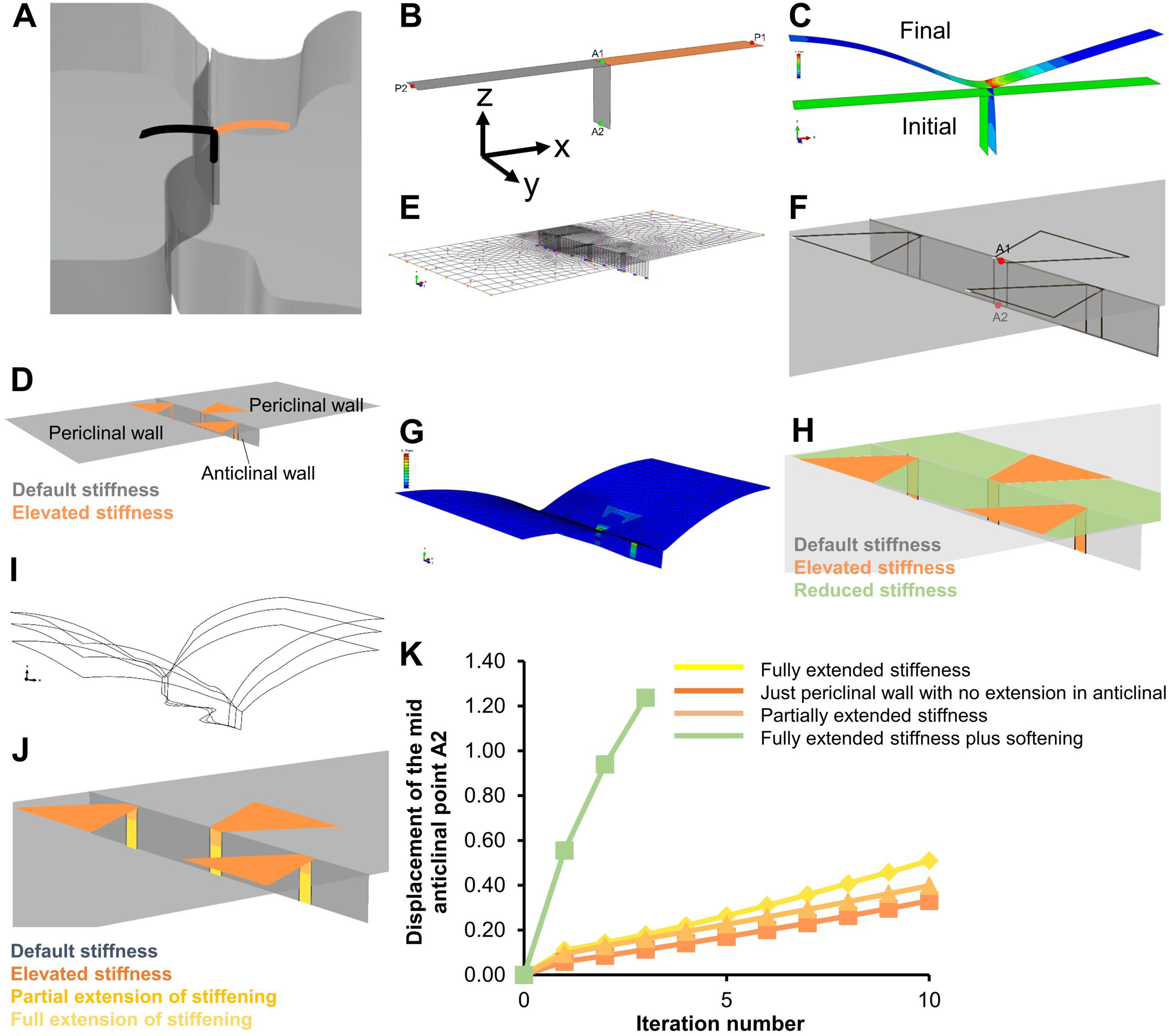
Model of cell wall segments containing stiff and soft regions deformed by turgor pressure. **A)** Illustration of the position a segment of two adjacent cells containing periclinal walls and the shared anticlinal wall extracted for simulations. The orange side refers to a higher stiffness or a lower extensibility. The deformation of this segment under turgor application is modeled using beam and shell approach. **B)** Shell model of wall segments. Pressure load is applied on lower surface of the periclinal walls, similar to loading in 2D beam model shown in Fig. S2A. **C)** Deformation in the shell model is similar to the beam model. **D)** Finite element model with alternate placement of stiffened regions on the periclinal and anticlinal walls of two adjacent cells. **E)** Turgor pressure is applied to inner surface of the periclinal walls (no pressure on the anticlinal wall as the acting forces cancel each other out). **F)** Position of fiducial markers used to monitor shape formation by measuring the displacement of the anticlinal wall. **G)** Initiation of undulations in the first iteration of the model. Heatmap represents von Mises stress. **H)** Finite element model with alternate placement of stiffened regions on the periclinal and anticlinal walls of two adjacent cells with cell wall softening in areas between the incipient necks. **I)** Evolution of undulations for three iterative load applications. **J)** Stiffening extensions on the anticlinal wall extending into different depths. **K)** Displacement of the anticlinal wall mid-point (A2) under different wall stiffening-softening scenarios reveals that fully extended stiffening leads to more pronounced undulation, but the effect is small compared to that of a softening in the periclinal wall. The stiff, soft and default regions were assigned Neo-Hookean elastic constants (*C*_10_) of 10, 0.05 and 0.5, respectively (see modeling procedures for more information).

In these simulations, the stiffness differential is implemented *ab initio*—prior to application of the turgor pressure (*ab initio* stiffening). Simulations reveal that it does not matter for the outcome whether the cells are under turgor (*cum tempore* stiffening) or relaxed (*ab initio* stiffening) when the stiffening of one of the periclinal cell walls is implemented (Figs. S2C-F). In a biological sense, this means that when a segment of the periclinal wall is reinforced, either by deposition of cellulose or by changes in other cell wall polysaccharides, a shift of the relative position of the anticlinal wall toward the cell with the stiffened periclinal wall can be expected upon the next step in development. This does not depend on whether or not the cell wall is under stress when the stiffening occurs.

### Turgor differential does not steer pavement cell morphogenesis in the presence of a stiffness differential in the periclinal wall

To verify the influence of turgor pressure on the formation of necks, differential turgor is applied to the two neighboring cells as detailed in the previous section. The results show that in all cases, regardless of the relative turgor pressures in the two cells, the anticlinal wall is displaced toward the cell with the stiffer periclinal wall (Figs. S3A-F) as it stretches less under tension compared to the softer side. In extreme conditions, when the pressure on the stiffened side approaches zero, the horizontal (in x direction) displacement of the anticlinal wall becomes negligible (Figs. S3E and F). This means that the mechanical and geometrical properties of the cell wall, and not a turgor pressure differential, dictate the evolution of lobes.

Further, it is observed that when the anticlinal wall is stiffer than the default value, i.e., the value of the non-stiffened periclinal wall, the displacement of the anticlinal wall (mid-points corresponding to points A1 and A2 in Fig. 3B) is enhanced as the anticlinal wall deforms less under the effect of the turgor differential (Figs. S3A, C, E). Taken together, the simulations suggest that turgor differential is not a requirement to form pavement cell undulations, but that cell wall mechanics seems to be dominant.

### The relation between stiffness differential and undulation is non-linear

To assess how different the stiffnesses between the lobe and neck on the opposing periclinal walls must be for an undulation to evolve, different ratios of stiffness between the stiffened and default periclinal walls are examined by monitoring the resultant displacement of the anticlinal wall. As the stiffness ratio increases from 1 to 5, the displacement of the anticlinal wall increases rapidly, but beyond this value the deformation plateaus (Fig. S4A). At this point, the stiffer side behaves as a rigid structure compared to the softer (lobe) side. Unless turgor pressure is increased, any additional stiffness in the stiffer wall does not result in a perceivable increase in horizontal (in x direction) displacement of the anticlinal wall.

We then tested whether subtle stiffness differentials would lead to undulations if load application is applied repeatedly. Iterative load application is accomplished by re-zeroing the stress developed in shell elements following each load application and by repeating the simulation, starting off with the last deformed shape. Running the simulation for 3 iterations shows that the resulting horizontal (in x direction) displacement of the anticlinal wall is negligible compared to the vertical (out of epidermis plane, in z direction) deformation (Fig. S4B). Therefore, when continued, the cells only balloon out of the plane, not forming discernable undulations. This demonstrates that repeated load application does not cause more pronounced wave formation if the stiffness difference is negligible to begin with and remains constant. A sufficiently large stiffness ratio between the two periclinal wall segments is required even when repeating the load application with stress relief (Fig. S4C).

### Reproducing an interlocking pattern of pavement cells requires alternate positioning of differential stiffness along the periclinal walls

The simulations thus far indicate that locally increased stiffness of a periclinal wall segment at one side of a cell-cell border can displace the relative position of the anticlinal wall in the presence of turgor. To verify whether the observed deformation in the periclinal segments and displacement of the anticlinal wall can reproduce multiple undulations arranged in series, finite element models of larger wall segments are developed to incorporate multiple, alternatingly placed regions of cell wall stiffening (Fig. 3D). To follow experimental observations on microtubule distribution, as will be shown in the experimental section, which presumably is translated to enhanced cellulose deposition, the stiffening regions on periclinal walls are extended in the depth of the anticlinal walls. For higher precision, the mesh of the finite element model is constructed with a higher resolution near the cell borders, and equal turgor pressures are applied in both cells (Fig. 3E). To assess shape deformation, horizontal (in x direction) displacement of the mid-point of the anticlinal wall (A2) is recorded in each simulation (Fig. 3F). The results of the simulations show that one iteration of pressure application causes a displacement of the anticlinal wall that simulates the initiatial expansion of lobes and necks (Fig. 3G). The distribution of stresses reveals that regions of higher stress correspond to the regions with higher cell wall stiffness. To attain more pronounced lobes and necks, the process of turgor application is repeated. After each iteration, the deformed geometry (output) is used as initial geometry (input) for the next iteration, and wall stresses are readjusted to zero. The subsequent simulation is performed with all other parameters kept constant (Fig. S4D). This model shows how continuous deformation of the cell wall with localized alternatingly placed stiffness on two sides of the anticlinal wall under turgor pressure can develop into lobes and necks.

### Softening of the cell wall promotes lobe outgrowth

It is thought that the delivery of new wall material and wall loosening enzymes on the lobe side of an undulation facilitate the expansion of lobe protrusions into the neighboring cells. The provision of this material is presumably mediated by arrays of actin microfilaments (Mathur and Hülskamp, 2002; Panteris and Galatis, 2005). To examine the effectiveness of wall softening in the generation of interdigitations, the finite element model is modified to allow for spatially confined softening of the cell wall. Similar to the stiffening, the softening is implemented either *ab initio* or *aum tempore*. The location of these softening regions is limited to the zones of the periclinal walls between the stiffened regions (green zones in Fig. 3H). As above, the triangular stiffened regions on periclinal walls are extended by narrow bands in the depth of the anticlinal wall. Simulations show that softening in the regions of incipient lobes facilitates extension of the wall interdigitation in xy plane while the ‘ballooning’ of the periclinal walls in the z direction is less pronounced compared to the simulations without the softening (Fig. 3I against S4D).

### Extension of stiffening in the depth of the anticlinal walls enhances but is not sufficient to derive the confined lobe growth

In the simulations above, the triangular stiffening regions in the periclinal walls are connected to extensions of stiffened bands in the anticlinal walls. To investigate the influence of anticlinal wall stiffness extensions, we allowed these bands to be only partially extended or absent and compared the displacement results with the full extension of the stiffened bands (Fig. 3J). The simulations indicate that in the presence of periclinal wall stiffenings, undulations can be generated without stiffening in the anticlinal wall, but the magnitude of the undulations is reduced in absence of the latter as measured by the relative displacement of point A2 (Fig. 3K).

Inversely, the presence of anticlinal wall stiffening in the absence of local stiffening in the periclinal walls produces no deformation under application of turgor pressure. This is consistent with the simulations made using the isolated anticlinal wall model (see Supplemental Notes 1 and 2). This simulation, therefore, further emphasizes the importance of accounting for the periclinal wall in the formation of interdigitations.

### Multi-cell simulation confirms that local stiffening and turgor-driven stretch of the cell wall can form interlocking pattern in the tissue context

To assess whether the concepts identified above hold for a multicellular context and to assess the emerging stresses, a multicell finite element model is developed by arrangement of multiple, identical whole cell modules with hexagonal geometry and alternatingly located stiffening and softening zones on the periclinal walls. The periclinal stiffening regions in inner and outer walls are connected by stiffened bands extending in the depth of the anticlinal walls (Fig. 4A). Upon application of turgor pressure, the cells form interlocking patterns (Fig. 4B). As in the previous sections, in this class of models a turgor pressure differential in adjacent cells is not necessary to generate the protrusions. The results indicate that highest stresses correspond to regions with higher stiffness (neck) (Figs. 4C). Interestingly, the model predicts that stress lines extend between adjacent necks and cross each other in the center of the periclinal cell wall (Figs. 4D and S4E).

**Fig. 4.**
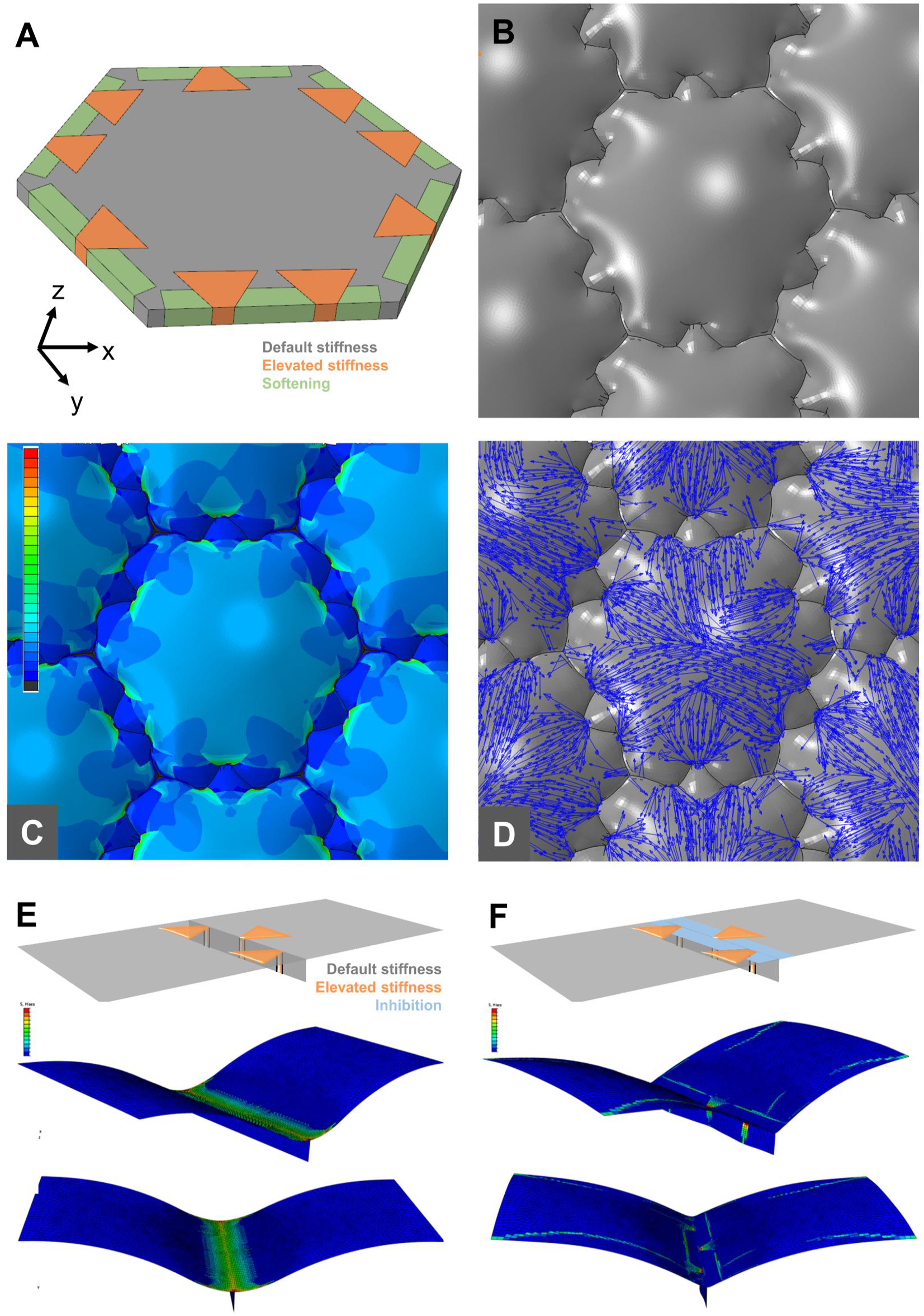
Shape and stress development in pavement cells based on inhomogeneity in cell wall stiffness. **A)** Whole cell model considering local, alternating stiffening and softening of the inner and outer periclinal walls as well as extension of the stiffened regions in the anticlinal walls. **B-D)** Simulation of turgor induced deformation and stress in a model composed of multiple single-cell units as shown in **A.** The turgor pressure was set equal in all cells. **B)** Shaping of undulations. **C)** Stress pattern generated by turgor application in the periclinal walls and at periphery of tricellular junctions. Heatmap represents maximum principal stresses. **D)** Orientation of stress lines in periclinal walls. **E-F)** Incorporation of positive stress-stiffening feedback mechanism that amplifies stiffness based on local stress conditions starting from minute stiffness differential (ratio 1.01) in periclinal and anticlinal walls. Deformation and stress pattern after hundred iterations of load application. Heatmap represents von Mises stress. **E)** Without inhibition mechanism stiffness differentials even out. **F)** Inhibition of stiffening at alternating locations allows for stiffness differentials to amplify and undulations to form.

To investigate the effect of the geometry resulting from the local stiffening and softening of the periclinal walls on the stress pattern in the cell wall, we used the output geometry of the previous simulation as a new input model, but we implemented isotropic and uniform material properties. The result of the subsequent load (turgor) application indicates that once the lobes and necks are formed, the geometry itself results in stress anisotropy with higher stresses in necks, even if their stiffness is the same as that of the lobe (Fig. S4F). This confirms the validity of our mechanical approach as the stress patterns predicted for lobes and necks developing *de novo* match the stress pattern of 3D pressurized cells presented above (Figs. 1E and F) as well as those in other studies (Sampathkumar et al., 2014). It further confirms that cell geometry is a crucial link in the morphogenetic loop (Kierzkowski and Routier-Kierzkowska, 2019).

### A positive mechanical feedback loop based on stress-induced stiffening and lateral inhibition can shape pavement cells

Previously, we showed that if the stiffness ratio between the opposing necks and lobes is not sufficiently large, iterating the simulation with repeated load application cannot produce discernable displacements on cell borders if the stiffness ratio remains constant. However, a living cell is dynamic; it can increase cell wall stiffness by adding cellulose. The deposition of cellulose microfibrils is guided by microtubules, and microtubules orient along stress fields (Landrein and Hamant, 2013; Uyttewaal et al., 2012). In the previous section, we observed that, even if the cell material possesses a uniform stiffness value in all regions after the lobes and necks are initiated, the shape itself generates higher stresses at the convex sides of the undulations on the periclinal walls. We, therefore, investigated the effect of stress-driven microtubule bundling on the cell shaping process if a feedback mechanism is implemented in which the cell wall is gradually stiffened in regions with locally elevated stress. This is implemented over several iterations starting with an infinitesimally small initial stiffness difference in the periclinal wall segments (Figs. 4E and F). After each iteration, the model is set to update local stiffness values based on the presence of local stress (Fig. S6A). For this purpose, a Python code was developed to extract the stresses present at each element of the structure at each iteration (output). Based on the local elemental stress, each element is assigned a new stiffness value calculated based on its previous stiffness value and the output stress resulting from the previous iteration (See Note 6, Supplemental Information).

The initial stiffness difference implemented for this test is 1%. The results indicate that the deformation upon the first iteration is relatively small and during subsequent iterations, the displacement of the lobe does not increase considerably but reaches a plateau with a negligible displacement (Fig. 4E). Stresses resulting from a single load application are not local enough to produce undulations through the feedback mechanism. Essentially, stresses are leveled out over the distances relevant here, and the feedback loop causes an increase in overall stiffness at the border of the two periclinal walls rather than sharpening the differences between neighboring lobe and neck regions.

We, therefore, tested whether adding an inhibitory mechanism could cause an augmentation in stiffness differential during feedback iterations. To implement this, feedback-driven stiffening is prevented in regions between the incipient necks, given as an input to the feedback model (Fig. 4F). Lobe formation is amplified by the feedback mechanism even when starting with very small stiffness differences (1%) between future necks and lobes (Figs. S6B and 4F). These results suggest that while a feedback mechanism starting from infinitesimally small stiffness differences can produce realistic results, it cannot be based on stress stiffening alone but needs to incorporate a mechanism that prevents stiffening at selected locations. It is worth noting that, in these simulations, we started from straight cell borders and did not incorporate the initial small waves that we postulate above due to cell wall buckling. When small initial waves are incorporated, the immediately appearing stress on the neck side might be sufficient to reduce the need for a stiffening inhibition mechanism.

## Experimental validation of *in silico* predictions

The models above predict that alternating changes in material properties of the cell wall at the site of undulations can produce an interlocking pattern in pavement cells. Changes in material properties suggest that variations in the biochemical composition of the cell wall are involved during the expansion of lobes. To experimentally validate the predictions made by the simulations, we studied the spatial distribution of two major cell wall components known to modulate the mechanical properties of primary plant cell walls: cellulose microfibrils and homogalacturonan (HG) pectin. This was performed in the epidermis of *Arabidopsis* cotyledons.

### Pectin is weakly esterified on the neck side

Our computational model predicts that the mechanical properties vary along the cell border on periclinal and anticlinal walls, but it does not specify which biochemical component of the cell wall is responsible for this effect. Pectin de-esterifìcation has been linked to changes in mechanical stiffness of the plant cell wall (Amsbury et al., 2016; Bou Dalier et al., 2018; Braybrook and Peaucelle, 2013; Carter et al., 2017; Chebli et al., 2012; Peaucelle et al., 2015). To investigate a possible involvement of changes in esterification of homogalacturonan (HG) pectin in spatial variation of mechanical properties of the pavement cell walls, we used COS^488^, an oligosaccharide probe conjugated with Alexa Fluor 488, recently synthesized and reported to be highly specific for de-esterified HG pectin (Mravec et al., 2014). Living cotyledons of wild-type *Arabidopsis thaìiana* taken from seedlings at 2-5 days after germination (DAG) were stained with COS^488^ and observed under the confocal laser scanning microscope (see experimental procedures and Note 7, Supplemental information). At this developmental stage, pavement cells have already acquired wavy shapes but continue to grow and generate new lobes (Zhang et al., 2011). We observed that COS^488^ signal exhibited variations along the profile of cell borders at the anticlinal walls (Fig. 5A). Maximum intensity projections of z-stacks were used to assess COS^488^ signal on both periclinal and anticlinal walls related to necks and lobes. The comparison of fluorescence intensity in maximum projections indicated a consistent, higher affinity of COS^488^ with the neck sides of undulations (p<0.0001, see Note 8, Supplemental Information, Figs. 5B and S7H). While in this study we mainly focused on pavement cells between 2-5 DAG, this association was not limited to mature cells or pronounced waves. Indeed, we observed that even at slight curvatures, a similar pattern of higher signal was associated with the neck side (Fig. 5C). Due to challenges of working with early-stage cells, we focused on day 2 after germination from here onwards. Even at this stage, cells are growing and new cells and new lobes continue to appear. Interestingly, COS^488^ staining also showed bright spots at cell junctions (Fig. 5A). We hypothesize that de-esterifìcation of pectin in the middle lamella may be employed at the junctions to enhance cell adhesion. The observation of bright signal at cell junctions is similar to the study by Carter et al. (2017) who also found a higher signal of COS^488^ at the poles of guard cells and associated this with polar stiffening in these cells. This consistency with other studies further confirmed the proper staining of samples for pectin using COS^488^ in our experiments.

A recent study has shown that propidium iodide, a general probe for the cell wall, has a higher affinity for weakly esterifìed pectin compared to the highly esterifìed form (Rounds et al., 2011). While propidium iodide cannot be reliably used as a specific marker of pectin compared to specific probes such as COS^488^, we nevertheless analyzed the signal intensity of samples stained with propidium iodide in a similar approach to the COS^488^ experiments. Similar to COS^488^ staining, the propidium iodide fluorescent signal intensity oscillated along the meandering borders of pavement cells (Fig. S5A). Consistently, maximum projections of z-stacks showed higher signal intensities on the neck sides of the undulations while at the lobe sides the signal was dim (Fig. 5D, S5B). Quantitative comparison of the signal intensity difference between neck and lobe pairs (n>100) of different cells (n=l 9) of several cotyledons (n=5), showed a significant statistical difference (p<0.0001). The same result was obtained when analyzing orthogonal views (Fig. 5D, bottom). This result is interesting as it reinforces the possibility that propidium iodide may indeed predominantly bind de-esterified pectin, as suggested by Rounds et al. (2011).

**Fig. 5.**
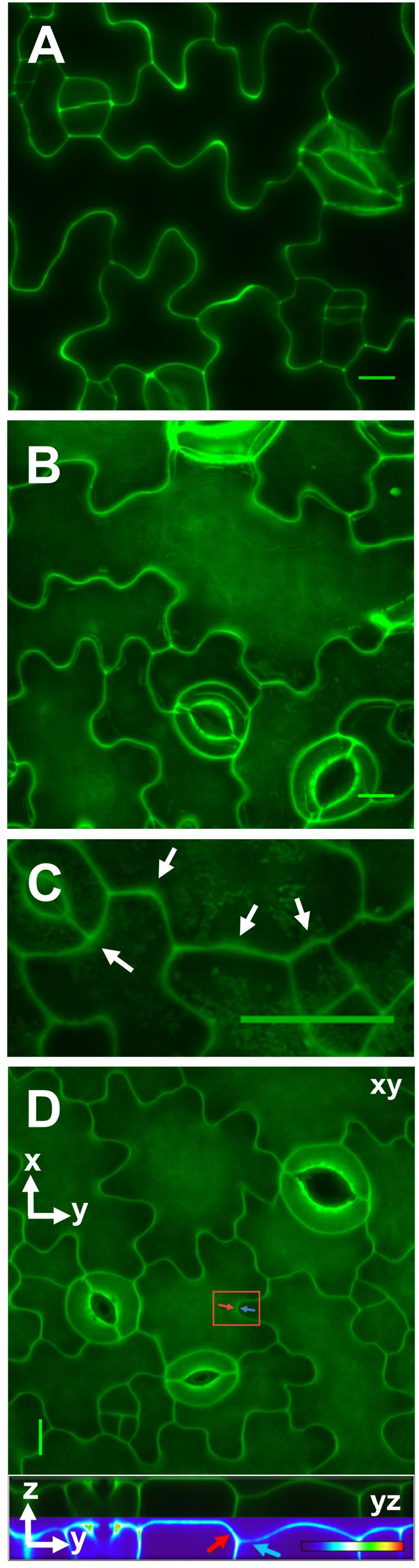
Staining for de-esterified HG pectin in *Arabidopsis thaliana* cotyledons sampled 3 to 5 DAG. De-esterified pectin when bridged with calcium increases cell wall stiffness. **A)** Single optical section through the center of the epidermal layer thickness stained with COS^488^, specific to weakly esterified pectin (Mravec et al., 2014), shows varying signal intensity along cell borders and high signal intensity at cell junctions. **B)** Maximum projection of z-stack with COS^488^ staining exhibits higher signal intensity at the neck side of the waves on periclinal walls (p<0.0001, for analysis see Fig. S7H and Note 8, Supplemental Information). **C)** Maximum projection of sample stained with COS^488^ shows that in smaller cells and along cell borders with barely developing bends, higher signal appears at the neck side of the periclinal wall similar to more pronounced waves as in **B. D)** (Top): maximum projection of z-stack labeled with propidium iodide and (bottom): orthogonal single yz optical slice through same z-stack, crossing the center of the marked lobe and neck pair in red rectangle (false color monochrome and heatmap). This orthogonal view was used to observe the cell wall at a pair of neck and lobe from the side. This view simplifies the illustration that the cell wall is brighter and that it bulges out on the neck side. In contrast, the periclinal wall on lobe side appears relatively dim (p<0.0001, see Fig. S5B, and Note 8, Supplemental Information) and flat. Comparison of fluorescence intensity between lobes and necks was carried out in circular regions of interest placed on opposite sides of undulations. Scale bars = 10 *μ*m (**A, B and D**) and 20 *μ*m (**C**).

To summarize, statistical analysis comparing the signal intensities between necks and lobes for both COS^488^, a specific marker for de-esterified pectin, and propidium iodide, showed higher signal intensities at the neck side and the differences with the corresponding lobes were significant (p<0.0001, see Note 8, Supplemental Information).

### Cellulose microfibrils and microtubules form organized bundles at the neck sides

The pattern of stiffening in the periclinal and anticlinal walls of pavement cells can be mediated through coordinated positioning of cellulose microfibrils which in turn is thought to be controlled by microtubules. We, therefore, investigated the orientation of microtubules in pavement cells and used cotyledons of GFP lines MAP4 and TUB6 of seedlings between 2-5 DAG. Visualizing the microtubules beneath the outer periclinal wall and along the anticlinal wall showed strong bundling of microtubules in association with necks (Figs. 6A-C and S7D, S5C). In lobes, microtubules could occasionally be observed, but their occurrence in this location was scarce compared to necks. It seemed that in central regions of cells, where bundles of microtubules arriving from multiple necks converged, the overall distribution of microtubules became random. A similar pattern was observed for stress lines in the multicell models of cell wave formation *de novo*, as discussed earlier (Figs. 4D and S4E).

**Fig. 6.**
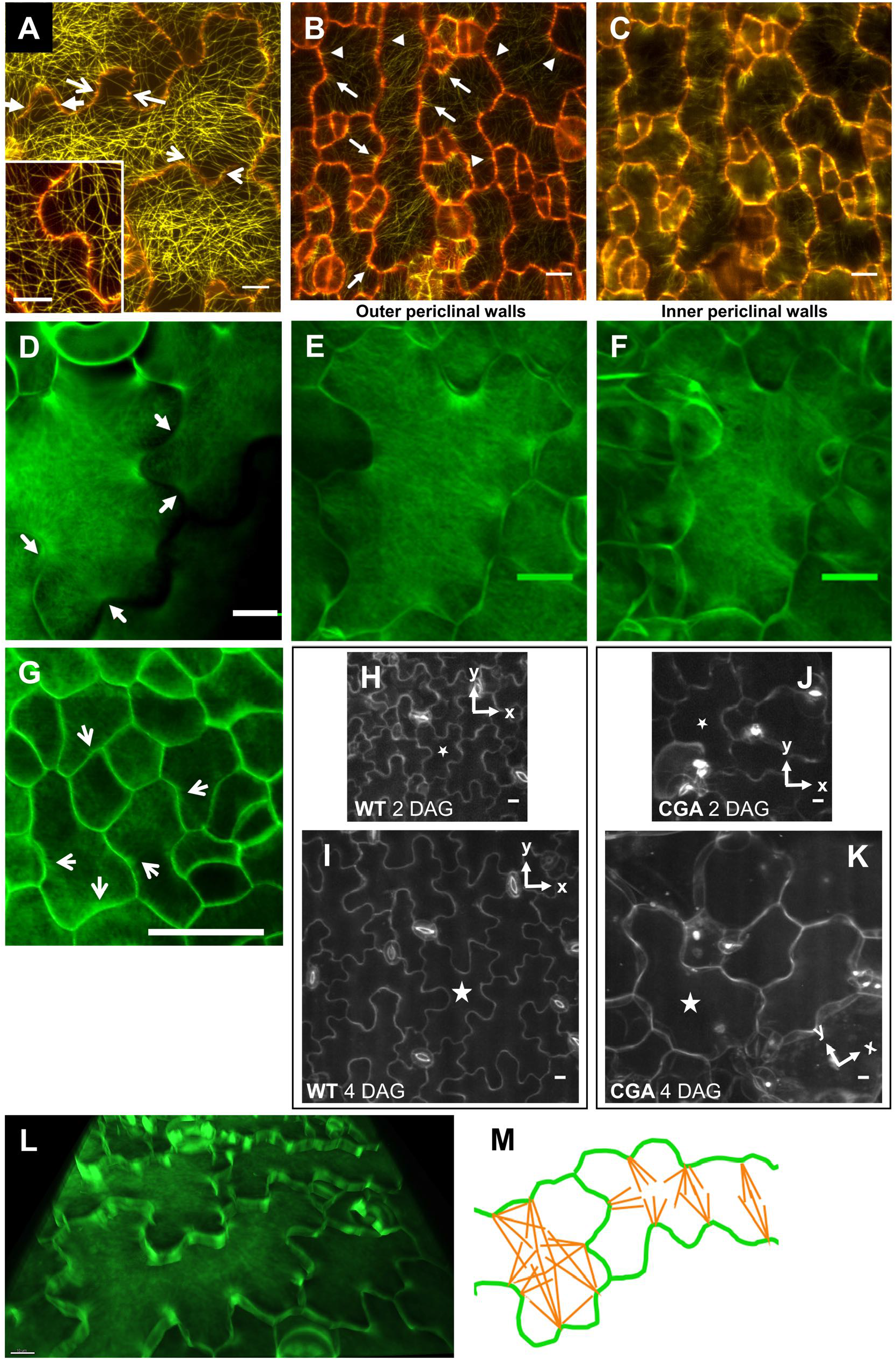
organization of microtubules (yellow pseudocolor) and cellulose microfibril (green pseudocolor). **A)** Microtubules labeled by GFP-MAP4 are abundant in association with the neck sides of undulations. Microtubules feature neck to neck connections forming circumferential hoops at the shank of lobes (pair of arrows, maximum projection). Inset shows higher magnification of a second sample. **B)** and **C)** Microtubules underlying the outer and inner periclinal walls, respectively, of the same pavement cells in the GFP-TUB6 line. Micrographs are maximum projections of the top **(B)** and bottom **(C)** halves of a z-stack corresponding to outer and inner periclinal walls, respectively. Microtubules were more abundant in neck sides of the undulations comparing signal intensity in selected circular regions on two sides of undulations (p<0.0001, see Fig. S7D and Note 10, Supplemental Information). In **A-C** a single optical section from the center of the z-stack was pseudocolored in red to mark cell borders. **D-G)** Calcofluor white staining for cellulose in pavement cells of *Arabidopsis thaliana* cotyledons at 3 to 5 DAG **(D, F)** and extracted from seed **(G).** For PFS staining of cellulose see Figs. S5D and E, Note 7, Supplemental Information. **D)** Cellulose microfibrils are concentrated at the necks from where they radiate into the periclinal wall (maximum projection, see also Figs. S5D-F and Movies S1 and S2). Cellulose microfibrils in outer **(E)** and inner **(F)** periclinal walls of the same cells. Micrographs show maximum projections of the top **(E)** and bottom **(F)** halves of a z-stack (see also S5H) using calcofluor white. **G)** Bundles of cellulose microfibrils in pavement cells of cotyledon extracted from the seed coat are associated with the necks of barely emerging curvatures. **H-K)** Time lapse of pavement cell morphogenesis of wild-type *Arabidopsis thaliana* cotyledons treated with DMSO (control) **(H, I)** and CGA **(J, K)** visualized at 2 and 4 DAG. Fluorescence is propidium iodide signal. While the DMSO-treated cells showed increase in surface area, perimeter and lobe number, the growing CGA-treated cells developed fewer lobes and necks (p<0.001, see Fig. S8 and Note 11, Supplemental Information). The coordinate systems indicate the orientation of the mounted samples. **L)** Oblique view of a reconstructed z-stack of pavement cells showing that cellulose microfibrils extend into the depth of the anticlinal wall at neck-lobe pairs (see Movies S1 and S2). **M)** Schematic representation of typical cellulose orientation depending on the aspect ratio of cells: in elongated cell regions (right), the microfibrils form a pattern predominantly perpendicular to the long axis. In cell regions with an aspect ratio close to one (left), bundles of cellulose are more centrifugally oriented (see also Fig. 4D). Scale bars = 10 *μ*m, 20 *μ*m (G).

We also used GFP-TUB6 line to confirm the observations (Fig. 6B, C and S7D). Similar to GFP-MAP4 pavement cells, necks were populated with microtubules in GFP-TUB6 cells (p<0.0001, Note 10, Supplemental Information). The fan-shaped orientation of microtubules was more evident in GFP-TUB6 results compared to the crowded arrays observed in GFP-MAP4. Interestingly, at some locations, microtubules appeared to form bundles at locations of the cell border that were only slightly bent. These sites were also associated with marked accumulations of microtubule label at the anticlinal walls (arrowheads marking the dotted microtubule on the red lines of cell borders in Fig. 6B). The inner periclinal walls showed a similar trend for the microtubules at locations of fully developed necks as well as for the relatively straight regions corresponding to arrowheads in Fig. 6B with a marked bundling of microtubules (Fig. 6C). We hypothesize that these regions may mark the locations for a strain/stress-driven positive feedback loop that result in fully-developed waves of lobes and necks. We hypothesize, as we will discuss later, that this focal polarization of microtubules is preceded by a mechanical isotropy breaking due to local cell wall buckling. Further, away from the tip of the lobes, bundles of microtubules radiating from two adjacent necks formed transverse bundles on the shank of the tube-like lobe (Figs. 6A). These arrays seemed to connect regions of greatest stress which the FE model predicts will develop in the necks. If this results in increased cellulose deposition at these locations, this stiffening would restrict the widening of lobes and instead allows for their elongation. Our finite element model showed that placement of stiffened necks and the shape of undulations produce stress fields that can eventually result in the orientation of microtubules that connect the opposing necks (Figs. 4D and S4E).

It is known that cellulose microfibril deposition is guided by cortical microtubules and, therefore, the most recently deposited microfibrils typically mirror the orientation or cortical microtubules. In the majority of the earlier studies on pavement cells, microtubule orientation has therefore been used as a proxy for cellulose-mediated wall stiffening, although direct evidence was rarely provided. The reason for this is mostly technical since fluorescent labeling of cellulose in living cells covered by a cuticle remains a challenge. To study the distribution and organization of cellulose microfibrils in pavement cells, we used calcofluor white and Pontamine Fast Scarlet 4B (PFS) (Anderson et al., 2010) to stain the living cotyledons (see Note 7, Supplemental Information). PFS is reported to be highly specific for cellulose and has been used in high-resolution imaging of cellulose in roots and onion scales (Anderson et al., 2010; Liesche et al., 2013).

In confocal images of pavement cells stained with PFS or calcofluor white, a notable concentration of cellulose microfibrils was observed in the neck regions from where they form a divergent fan-shaped configuration in the outer periclinal walls (Figs. 6D for calcofluor staining, S5D for PFS, Movies S1 and S2). Statistical analysis revealed a significant difference in PFS signal between necks and lobes (p<0.0001, refer to Note 9, Supplemental Information). Tips of lobes displayed considerably weaker signal and generally seemed devoid of organized microfibrils. Similar to the organization of microtubules, microfibrils appeared to connect neighboring necks (Fig. 6D arrows. S5D-E). Similar to the organization of microtubules, in most pavement cells of seedlings between 2-5 DAG, the orientation of cellulose microfibrils in outer periclinal walls away from the undulations were mostly random in mid-regions exhibiting lower aspect ratios or dominantly transverse in narrow cell regions that were axially elongated (Figs. 6D, M and S5D-F). The association of cellulose bundles with necks was not limited to pavement cells that displayed pronounced lobes. We observed that even in pavement cells of cotyledons acquired prior to germination by removing the seed coat, cell shape curvatures were already associated with radiating cellulose bundles (Fig. 6G).

To investigate the mechanical impact of the observed spatial distribution of cellulose microfibrils on wavy cell shape development, we experimentally reduced cellulose crystallinity and monitored the shape of the same cells in a time-lapse study. Reducing cellulose crystallinity is suggested to correlate with altered cell shapes and occurrence of a swollen phenotype (Aouar et al., 2010; Fujita et al., 2013). We grew *Arabidopsis* seedlings in presence of CGA 325 ‘615 (CGA), shown to reduce cellulose crystallinity (Crowell et al., 2009; Peng et al., 2001), and monitored shape parameters over 4 DAG. At 2 DAG, cells in samples grown in media containing CGA (treated cell hereafter) and the control cells had the same perimeter and area (see Note 11, and Fig. S8, Supplemental Information). However, lobe formation in treated cells was significantly altered. Following the same cells over a period of 4 first days after germination showed that control cells had an average of 9.1 (± 0.7) lobes, whereas treated cells only developed 3.3 (±0.5) at 2 DAG. Over the two subsequent days (at 4 DAG), control cells formed 6-7 additional lobes, whereas CGA treated cells only added 1-2 lobes despite similar increase in cell size (p<0.001, see Note 11, Supplemental Information for analysis and Fig. S8). These results indicate that interference with cellulose caused a significant reduction in lobe formation from early stages of epidermal cell development (Fig. 6H-K). This corroborates the significance of cellulose in development of wavy cell borders by restriction of expansion in the neck regions. Also, this further emphasizes that the present model is not limited to cells that are already highly wavy and that this mechanism is essential also lobe formation at earlier stages of development. These observations support the predictions made by the finite element model incorporating alternate placement of stiffness on periclinal walls.

Acquiring images of the cellulose orientation in the inner periclinal walls proved challenging due to the thickness of the samples and the fact that the signal from the inner periclinal walls was difficult to distinguish from the signal deriving from the directly adjacent mesophyll cell walls. However, in some cases, we succeeded and observed that the cellulose enrichment and orientation on the inner periclinal wall seemed to mirror the outer periclinal wall (Figs. 6E, F and S5H). This prompted us to analyze the cellulose status of the anticlinal walls since they might serve as mechanical links between the inner and outer faces of pavement cells.

### Cellulose microfibrils and microtubules form bundles at the undulations that extend in depth of the anticlinal walls coupling the inner and outer walls

3D reconstruction of confocal images of cotyledons stained with calcofluor white revealed that in neck regions cellulose extends down the anticlinal walls (Fig. 6L, Movie S2). This can also be observed by orthogonally slicing the z-stack and acquiring maximum x or y projections for narrow segments (Figs. S5F1 and F2). As the finite element simulations show, the anticlinal extension of cellulose microfibrils at the undulation can act as a lever, increasing the magnitude of undulations.

Regardless of the mechanical contribution of the anticlinal reinforced bands in the development of undulations, they might form due to and transmit the mechanical signal from the inner periclinal wall up to the outer periclinal wall or the other way around. To visualize this, we used the finite element model with the extension of stiffening in the depth of the anticlinal wall but without stiffened regions on the periclinal wall (Fig. 7A). Although this configuration does not generate any undulation in the anticlinal wall, the stress can be partly transmitted to the periclinal wall at its junction with the anticlinal bands. Such a transmission of stress has the potential to mark the position, attract further bundling of cortical microtubules in a positive feedback loop, and lead to deposition of cellulose under the outer periclinal wall. Since the stress due to local stiffening of the inner periclinal wall can be transmitted to the anticlinal walls (Fig. 7B), these bands of anticlinal stiffening can originate from the stress generated from the inner periclinal wall. This may explain how, if one of the periclinal walls develops a particular pattern of anisotropy, the other wall is triggered to mirror this through the microtubule-mediated addition of cellulose microfibrils. The stress pattern would be propagated from one periclinal wall to the other by way of the anticlinal walls.

**Fig. 7.**
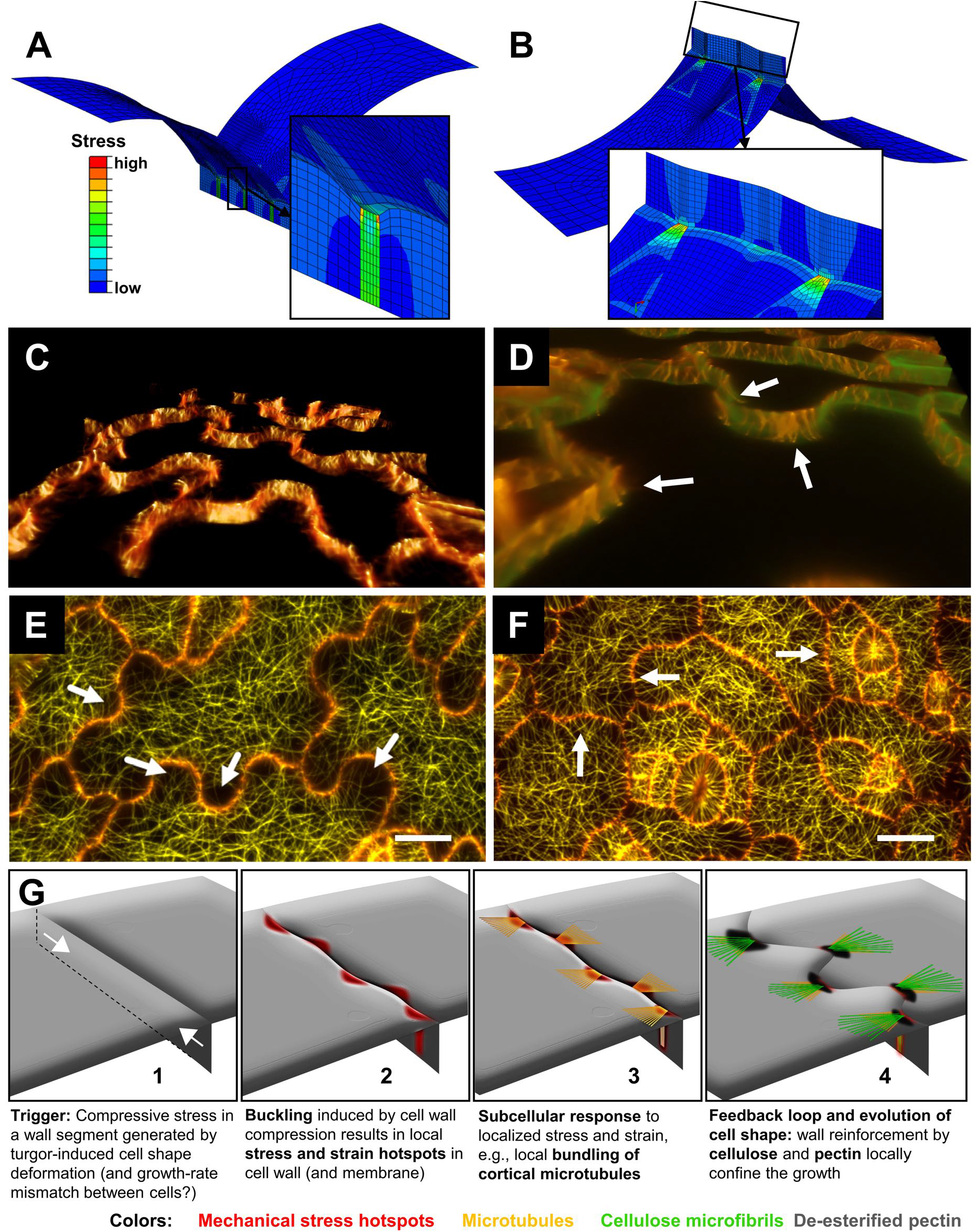
Patterns of mechanical stress and microtubule orientation. FE models containing a segment of anticlinal and periclinal walls under turgor pressure show that local stress raised in either **A)** anticlinal or, **B)** in the periclinal walls is transmitted to the respective connecting wall forming a stress coupling between the inner and outer periclinal and the connected anticlinal cell walls. In these cases, elevated stress resulted from a band of stiff inhomogeneity embedded in each of these walls. **C)** Oblique view of a z-stack 3D reconstruction of pavement cells from *Arabidopsis thaliana* GFP-MAP4 line excluding periclinal walls demonstrating cortical anticlinal microtubules. **D)** Dual channel image of propidium iodide (green) and GFP-MAP4 (orange) showing that cortical anticlinal microtubules were more abundant at the neck side of the undulations (p<0.0001, determined by counting the number of microtubules on each side, see Note 10 Supplemental Information and Movie S4). **E** and **F)** Cortical periclinal microtubules visualized in cotyledons of the *Arabidopsis thaliana* GFP-MAP4 line at 4 **(E)** and 1 DAG **(F)**. At both stages, microtubule density at opposite sides of corresponding lobe-neck pairs is different with higher density at the neck side. However, this difference is less pronounced at earlier developmental stages (white arrows in **F)**. At 4 DAG, microtubules appear to be bundled in necks while they are scarce in lobes. Occasionally microtubule bundles reach the tips of a lobe. The images do not correspond to the same cotyledon. Maximum projections of z-stacks. Scale bars = 10 μm. **G)** Schematic of the proposed multi-step process initiating pavement cell morphogenesis. **G-1)**. Compressive stress in anticlinal walls arises due to internal pressure bulging out periclinal walls. **G-2)** Stress inhomogeneities arisen due to buckling create stress hotspots that trigger local bundling of microtubules **(G-3)**, alterations in pectin esterification and reinforcement of the cell wall by cellulose deposition. Feedback mediated cellulose deposition leads to further expansion of lobes **(G-4)**.

Interestingly, after removal of the optical slices pertaining to periclinal walls to reveal the anticlinal wall, 3D reconstruction of the confocal stack shows that cortical microtubules assumed a preferentially vertical (z direction) orientation on the anticlinal walls (Figs. 7C and D, Movie S4). Since microtubules are thought to orient predominantly in direction of maximal stress, the vertical orientation corroborates the presence of a stress pattern oriented perpendicular to the plane of the leaf, as predicted by our finite element model (Fig. 2B). This suggests that putative stresses due to “tissue-level” forces that would be inherently parallel to the leaf surface in the anticlinal walls, even if present in leaf epidermal cells, are overridden by substantial cell-level turgor-generated stresses. Importantly, we observed that distribution of microtubules on the anticlinal walls was non-uniform and the microtubules were enriched near the locations of necks. Projecting the anticlinal microtubule signal from a 3D z-stack onto a 2D plane can render the precise localization of microtubules challenging. This is because microtubule-enriched regions placed alternatingly on two sides of the thin anticlinal wall can become impossible to associate with one side or the other. 3D visualization of microtubules (Movies S3-S5) and specifically using dual channel 3D reconstructions (with propidium iodide marking the cell wall, Movie S4) was helpful to determine microtubule localization relative to the adjacent anticlinal wall, particularly so in case of early-stage bends (shallow bends marked in Movie S5). Our analysis of microtubule density between lobes and necks demonstrated that in the majority of cases, microtubules were enriched on the neck sides of the anticlinal walls, coinciding with microtubule bundling under periclinal wall necks (p<0.0001, see Figs. S7F1 and F2, Movie S5 and Note 10, Supplemental Information) Thus, microtubule association with necks is not limited to periclinal walls, but also occurs at anticlinal walls.

### Cell wall buckling precedes microtubule polarization

An important observation of this study resulted from comparing the distribution of microtubules at different stages of lobe development. As discussed in the previous section, using GFP tagged microtubule lines, we noted that microtubules appeared to be bundled in necks in regions with an already established curvature, while they were scarce in lobes and only occasionally did cortical microtubule arrays seem to stretch into the tip of lobes (Fig. 7E). In smaller, less developed cells or regions of developed cells with walls with smaller bends, however, the difference in microtubule density on two sides of the wall between lobes and necks appeared to be significantly less pronounced compared to later stages (Fig. 7F). Our analysis plotting the difference in microtubule numbers across the wall versus the degree of wall curvature (as a measure of lobe progression) revealed a moderate correlation (r=0.5, see Figs. S7E, G and Note 10, Supplemental Information). This is consistent with the notion proposed here, that microtubule bundling and consequent wall reinforcement can amplify the amplitude of undulations in a positive feedback loop following an initial buckling event, rather than being the initial trigger.

## Discussion

Unlike animal cells, in plant cells actin-myosin based actuators are not directly involved in generating the forces required to move or alter cell shape. In this study, we show that complex cell shaping in plants is enabled by localized mechanical modification of the cell wall. Interlocking leaf epidermal cells are common in many plants and have attracted considerable attention over the past decades. Still, neither the events underlying the genesis of these shapes nor their purpose is well understood (Jacques et al., 2014; Vőfély et al., 2019). The analysis of the underlying molecular pathways and the characterization of the roles of cytoskeletal activities and cell wall biosynthesis have presented a formidable challenge to biologists and modelers (Armour et al., 2015; Belteton et al., 2017; Bidhendi and Geitmann, 2019; Fu et al., 2005; Majda et al., 2017; Panteris and Galatis, 2005; Sampathkumar et al., 2014; Sapala et al., 2018; Sotiriou et al., 2018; Xu et al., 2010; Zhang et al., 2011). Until now, the focus of research has predominantly been on the localization of the cytoskeletal components and the evaluation of their contribution to cellular “lobiness” (Armour et al., 2015; Belteton et al., 2017; Fu et al., 2005), whereas experimental evidence for events pertaining to the cell wall composition and a mechanical model explaining the genesis and development of the interlocking cellular protrusions have remained scarce (Majda et al., 2017). A study by Sampathkumar et al. (2014) analyzed the correlation between cell wall stress and microtubule organization in pavement cells, albeit *post facto*, i.e. in cells that were already wavy. This study crucially consolidated the existing knowledge and linked microtubule organization, cell shape and mechanical stress, but the origin of complex cell shapes was not addressed. Here, we focused on events leading to wavy cell shapes *de novo* and how these shapes are further developed to eventually yield an interlocking cell morphology. Based on our experimental data and mechanical modeling we propose a new paradigm in which the compressive stresses in cell wall segments—due to turgor pressure and cell shape—result in cell wall buckling. Anisotropy in strains/stresses arising from buckling at cell borders form hotspots invoking subcellular responses such as local bundling of microtubules. Such events result in the reinforcement of the cell wall by cellulose deposition and changes in pectin network, reinforcing the neck side of the small waves formed. The reinforced necks form regions of limited growth while lobes elongate anisotropically. Further, stress inhomogeneity arising from reinforcements form a feedback loop establishing lobe elongation and the interlocking morphology of pavement cells. Crucially, our simulations and microscopic observations of microtubule arrangement suggest that mechanical events can act as a cell-scale morphogen that precedes biological or cellular responses such as changes in cytoskeletal dynamics.

Our initial simulations based on isolated anticlinal walls (see Notes 1 and 2, Supplemental Information) indicate that the mechanics of epidermal cells and, specifically, the stress state in the cell wall cannot be adequately addressed without taking the periclinal walls into account. Separately, we showed that a stretch-based model of an isolated anticlinal wall as proposed by Majda et al. (2017) can only produce near imperceptible bends. These bends are virtually eliminated when periclinal walls are added, and the approach raises a number of inconsistencies elaborated in Bidhendi and Geitmann (2019), thus confirming that isolating a geometrical feature may not always be an admissible modeling simplification for pavement cells.

Simulations including the periclinal walls in turgid cells, on the other hand, generate crucial and non-intuitive information. When turgor pressure is applied to all walls in a complete 3D model, with the cell dimensions typical for epidermal cells of *Arabidopsis*, the periclinal walls bulge out and the anticlinal walls are pulled inward reducing the inplane projected cell surface (Fig. 2D). This occurs despite the exertion of the same pressure on the anticlinal walls. The reason for this is that due to the difference in surface areas, the combined forces resulting from the turgor pressure on the periclinal walls are higher than those acting on the anticlinal wall surface. These simulations also imply that for a cell to grow in-plane, selective and isotropic softening of the anticlinal walls is not sufficient and changes in mechanical properties of periclinal walls are required. Another important conclusion obtained from 3D cell models is the finding that anticlinal walls experience high stresses vertically (in the z direction) compared to the in-plane directions (Figs. 2B and C). This stress pattern matches the net orientation of cortical microtubules underlying the anticlinal walls. The presence of vertical microtubules suggests that anticlinal walls become reinforced anisotropically by cellulose microfibrils starting at the very early stages, upon formation and fusion of the cell plate. This anisotropic reinforcement enables the anticlinal walls to grow in their longitudinal direction, perpendicular to the net orientation of microfibrils, allowing the growth of pavement cells in-plane while preventing their out-of-plane expansion (Figs. S7A-C, Movies S3 and S4).

Our simulations indicate that the formation of alternately located, non-yielding (stiff) regions on periclinal walls in a growing cell suffices to develop necks and lobes. We show that this alternating pattern of stiffness distribution is consistent with the pattern of cellulose deposition and de-esterification of pectin. The stiff regions form the indentations or necks. Our simulations show that once lobes and necks are initiated, the geometry itself will cause the appearance of elevated stress at the neck side (Figs. 1E, F, 4C, D and S4E, F) even if isotropic material properties are used. The locations of elevated stress and the anisotropic stress pattern obtained through static simulation closely match the experimental observations of cortical microtubule bundling in necks and their orientation (Figs. 4D, 6A-C, 7D and E). This model can mechanically explain why pharmacological interference with cortical microtubule bundling has the potential to produce pavement cell phenotypes with reduced waviness (Armour et al., 2015; Fu et al., 2005; Mathur, 2004). However, the fact that in all cases of pharmacological interference with actin and microtubules, there remains at least a slight degree of border waviness is a matter that can be explained by the spontaneous buckling of the walls, as will be discussed later.

In the positive mechanical feedback loop between local wall stress and stiffness, we demonstrate that developing lobes and necks can initiate from infinitesimally small stiffness differentials if a stress-stiffening mechanism with lateral inhibition is incorporated. The existence of such a lateral inhibition mechanism seems to find an immediate biological equivalent in the proposed antagonism and competition between microtubule and actin regulation pathways (Chen et al., 2015; Fu et al., 2005; Fu et al., 2002; Xu et al., 2010). Since microtubules are associated with guided deposition of cellulose, and actin filaments are thought to deliver new, and presumably softer, wall material, such a scenario would cause lateral inhibition of stiffening as predicted by our model. The antagonistic pathways and, specifically, the exclusivity of actin to lobes are not uncontested, however (Armour et al., 2015; Belteton et al., 2017). For instance, Armour et al. (2015) did not observe any obvious association between lobes and actin microfilaments at least in the early stages of pavement cell shape development. Therefore, the roles and the competition between the cytoskeletal elements in driving the progression of cellular protrusions warrants further investigation.

Further in the shank of lobes, bundles of microtubules radiating from two neighboring necks form transverse connections crossing the shank of the tube-shaped lobe (Figs. 6A-C, and S7D). This structural pattern matches the stress pattern predicted by our finite element model (Fig. 4D and S4E). This pattern of cortical microtubule arrays is clearly translated into cellulose orientation at these locations (Figs. 6D, S5D-F) presumably restricting further widening of lobes while promoting anisotropic expansion in length of lobes. These observations and predictions are consistent with anisotropic growth patterns in pavement cells reported by Elsner et al. (2018).

Staining the cotyledons with COS^488^ shows pectin in pavement cells to be weakly esterifìed at the neck sides of undulations. De-esterified pectin can enhance the mechanical stiffness of the pectic network when cross-linked by calcium. Therefore, changes in the pectin status in the neck region could have a direct consequence for local cell wall stiffness in periclinal walls. Variations in the esterification status of pectin have been implied in other plant tissues related to the anisotropic growth of cells, cell shape deformations and organogenesis (Bidhendi and Geitmann, 2016; Carter et al., 2017; Majda et al., 2017; Palin and Geitmann, 2012; Peaucelle et al., 2015; Wolf et al., 2009). It is possible that de-esterification of pectin in necks locally decreases cell wall deformability by preventing slippage and separation in the xyloglucan-cellulose network thus locally impeding cell wall growth (Abasolo et al., 2009).

Staining cotyledons with calcofluor white and PFS revealed bundles of cellulose microfibrils radiating from the neck side of the undulations confirming a link between mechanical stress, microtubules and biochemical makeup of the cell wall. Accumulation and bundling of cellulose microfibrils in these regions can be interpreted as locally elevated stiffness of the cell wall. This is due to a considerable tensile stiffness of cellulose bundles compared to other cell wall polysaccharides. The tips of lobes on the periclinal walls showed considerably less fluorescent signal and generally seemed devoid of organized cellulose microfibrils. Similar to the organization of microtubules, microfibrils seemed to connect neighboring or opposite necks (Fig. 6D). In some observations, the neck-neck connections in the inner periclinal walls appeared to be more pronounced, and bundles of cellulose seemed to be more tightly packed compared to their corresponding outer periclinal wall (Figs. 6E and F and S5H). Therefore, the inner periclinal wall may be more determinant in dictating the augmentation of undulations while the outer periclinal wall may loosely mimic the cellulose pattern of the inner wall. Such harmonization is possible by stress and strain coupling through the formation of anticlinal strips of cellulose reinforcement as we showed through modeling (Figs. 7A, B). Alternatively, the observed stronger neck to neck associations and tighter bundling in the inner periclinal walls may reflect a temporal order of events with the development of anisotropic reinforcement in the inner wall of a cell preceding that in the outer periclinal wall.

Armour et al. (2015) reported that the regions of the anticlinal wall where microtubules form the cortical anticlinal bundles mark the position of incipient necks. However, close analysis of their micrographs suggests that even the earliest evidence of microtubules focally marking a region of the anticlinal wall is associated with an already existing local change of curvature, even if that curvature is not very pronounced. Therefore, the available studies cannot indicate whether there was a change in the cell wall composition or stress that caused bundling of the microtubules in those regions or whether it was the microtubule accumulation that predicted the site of future lobes. Importantly, in our study we noted that the change in microtubule density on the two sides of an undulation is a gradual process and the difference on the opposing sides is imperceptible for slight wall curvatures (Fig. 7E and F). We suggest that the initial difference in microtubule density is a response to a difference in mechanical stress on two sides of an undulation. Our new hypothesis considers this stress difference to result from an event that forms slight waves in the isotropic cell walls through buckling and that precedes the subcellular polarization or cell wall anisotropy. Buckling is a mechanical instability that occurs when structures lose their axial load-bearing capacity under compressive stress states resulting in formation of bends. Here we suggest this mechanical phenomenon to act at subcellular level as a morphogenetic step and a polarizer leading to the spontaneous and stochastic formation of local stress hotspots (Fig. 7G) and eventually to mechanical anisotropy and the formation of complex cell shapes. We provide a proof of concept for the notion that buckling in the walls can simply result from compressive stresses arising from cell geometry and turgor (Fig. 2). Localized stress can then trigger microtubule bundling and, as explained earlier, acting in a positive feedback loop, can result in cell wall reinforcement through local cellulose reinforcement. A schematic of the proposed morphogenetic steps summarizes the events in Fig. 7G. This concept does not exclude that independently, and compounding this effect, external compressive stresses may arise due to conditions such as variations in growth rates between neighboring cells. Indeed Armour et al. (2015) and Elsner et al. (2012) demonstrated that cell wall growth rates can vary considerably between neighboring pavement cells as well as between different regions of an individual cell. Further, buckling of a structure depends not only on the magnitude of the axial compressive loads but also on the geometrical aspects of that structure, e.g. wall thickness. Based on this, we propose that the difference in dimension of epidermal cells is one of the factors that influences how resilient epidermal pavement cells are against buckling, thus creating the great variation of waviness observed among pavement cells of different species (Vőfély et al., 2019).

Last but not least, our proposed model that is based on buckling as a mechanical event preceding cytoskeletal and cell wall polarization also provides a conceivable explanation to a phenomenon that is subtle and hence typically neglected: the presence of residual undulations in in specimens subjected to pharmacological treatments or mutations. Interference with the regulators of microtubule functioning such as brassinosteroids, regulators of ROPs such as RhoGDIs, mutations in microtubule-associated or microtubule-severing proteins such as IQD, CLASP or KATANIN all considerably decrease but in no case do they completely eliminate cell border undulations (Akita et al., 2015; Ambrose et al., 2011; Fu et al., 2005; Han et al., 2015; Kirik et al., 2007; Kotzer and Wasteneys, 2006; Liang et al., 2018; Lin et al., 2013; Liu et al., 2018; Mitra et al., 2018; Möller et al., 2017; Wu et al., 2013). The same applies to mutations or treatments affecting the actin cytoskeleton (Le et al., 2006; Rosero et al., 2016) or cellulose synthesis or crystallinity such as in *any1* (Fujita et al., 2013; Ivakov and Persson, 2013) or our CGA treatments. In other words, neither interference with cytoskeletal elements nor alteration of the wall composition has shown to result in complete loss of waviness at cell borders. Such events can always be explained to be due to the existence of redundancies and compensatory mechanisms, but our model raises a different hypothesis: while microtubules and subsequent cellulose reinforcement of the cell wall are crucial in pavement cell development and morphogenesis, they do not initiate cell border undulations but establish and enhance them.

## Experimental Procedures

### Plant growth conditions

In addition to the information below, for detailed experimental, computational modeling and statistical analysis procedures, refer to the Notes in Supplemental Information.

### Plant growth conditions

*Arabidopsis thaliana* ecotype Col-0 seeds were germinated in sterile Petri plates containing 1X MS (Murashige and Skoog, 1962) media with 1% sucrose and 0.8% plant agar under long-day lighting condition. The seeds for GFP-expressing line GFP-TUB6 (Nakamura et al., 2004) was obtained from Arabidopsis Biological Resource Center, under stock number CS6550. The GFP-MAP4 line (Marc et al., 1998) was also used to verify the observations with GFP-TUB6.

### Staining

Staining procedures were carried out mostly in dark condition. For pectin, seedlings were stained with either propidium iodide or COS^488^. COS^488^ was generously provided by Dr. William George Tycho Willats (University of Copenhagen). For visualizing cellulose, calcofluor white and PFS, a dye with a high affinity to cellulose, were used. Calcofluor was used at a concentration of 2 mg/mL in ddH_2_O. PFS staining was carried out with a 14 mg/mL solution of PFS in PBS buffer (Na_2_HPO_4_ 3.2 mM, KH_2_PO_4_ 0.5 mM, NaCl 135 mM, KCl 1.3 mM, pH 7.3).

### CGA treatment and time-lapse study of pavement cell growth

Cotyledon samples from *Arabidopsis* were treated with the herbicide CGA (CGA 325’615) at 0.9 nM prepared from a 10 μM stock solution dissolved in DMSO. CGA is suggested to inhibit the synthesis of and reduce the cell wall content of crystalline cellulose (Peng et al., 2001).The same concentration of DMSO (v/v) was used for the control experiment. The solutions were added to the ½ x MS (Murashige and Skoog, 1962) growth media. The samples were labeled with propidium iodide (0.01 mg/ml) for 20 min, followed by three washes with distilled water before observation. Propidium iodide labeling was applied at each time point prior to observation. Samples were mounted between slide and coverslip at each image acquisition. After each image acquisition, samples were placed immediately back to the in vitro growth chamber. At each time point, the same cells were located and traced. The adaxial side of the wild-type *Arabidopsis* was chosen for the study. For statistical analysis, for each time-point, 50-70 cells were studied from 10-12 seedlings.

### Fluorescence microscopy

Fluorescence microscopy was carried out on a Zeiss LSM 510 META confocal laser scanning microscope using a Plan Apochromat 63x oil immersion objective with numerical aperture of 1.40. For propidium iodide and PFS, excitation wavelength of 532 nm and bandpass emission filter of 550-615 nm was used. For COS^488^, 489 nm laser with bandpass filter of 550-615 nm was used. For calcofluor white, excitation wavelength of 405 nm in META mode and bandpass filter of 420-480 nm was used. For GFP lines, either excitation wavelength of 489 nm with emission bandpass of 500-525 nm, or in META mode, the argon laser of 488 nm with bandpass filter of 505-550 nm were used. For time-lapse imaging of CGA-treated and control samples, LSM 5 LIVE was used with 532 nm laser with an emission filter 590-625 nm.

### Image analysis and 3D reconstruction software

Analysis of fluorescence intensity was performed on maximum projections of z-stacks using ImageJ (Schneider et al., 2012). 3D reconstruction of confocal z-stacks was carried out using either Amira 5.6.0 (FEI, Visualization Science Group) or Bitplane Imaris 7.5.2 (Bitplane A.G.). Supplemental Movies were created using the volumetric rendering function of Imaris software on z-stacks acquired from the confocal microscope. For measurements and statistical analysis pertaining to cell wall components or microtubules refer to the Notes in Supplemental Information.

### Modeling Finite element simulations

Abaqus 6.14-2 finite element package was used for the creation of the geometries, meshing and post-processing. Abaqus/Standard solver was used for quasi-static finite element simulations (see Notes 3 and 4 in Supplemental Information on modeling procedures and Abaqus, 2014 user manual for more details).

### Feedback loop

To implement a feedback loop, a Python script was developed to read and write in the finite element model. After each iteration, the code extracts the deformed geometry from the Abaqus database and reads the stresses for each element. If a specific element has a stress higher than a threshold and does not belong to a list of stiffening-inhibition zone, the new value of stiffness for that element is updated in a stress-stiffening paradigm (see Note 6, Supplemental Information)..

## Author contributions

A.J.B. and A.G. designed the study, A.J.B. performed the modeling and the experiments. B.A provided the CGA treatment results. A.J.B., F.P.G, and A.G. analyzed the data, A.J.B. and A.G. wrote the manuscript.

## Acknowledgments

This project was supported by a Discovery grant from the Natural Sciences and Engineering Research Council of Canada (NSERC) and by the Canada Research Chair Program. We would like to thank Dr. William George Tycho Willats and Dr. Jozef Mravec from the University of Copenhagen for kindly providing us with COS^488^ for pectin staining. We would like to thank those colleagues who have provided constructive comments on previous versions of this manuscript as these helped us to improve this work.

## Supplemental Movies

**Movie S1**. 3D reconstruction of confocal z-stack demonstrating pavement cells of *Arabidopsis* cotyledon stained with PFS for cellulose. Locations of necks (indentations) on periclinal walls are clearly associated with a markedly higher signal. Related to Figs. S5D and E.

**Movie S2**. 3D reconstruction of confocal z-stack demonstrating pavement cells of *Arabidopsis* cotyledon stained with calcofluor white for cellulose. Locations of necks (indentations) on periclinal walls are clearly associated with a markedly higher signal. The cellulose enriched zones are present on both anticlinal and periclinal walls. Related to Figs. 6D-G, L and S5F and H.

**Movie S3**. 3D reconstruction of confocal z-stack demonstrating microtubules in pavement cells of *Arabidopsis* cotyledon from GFP-TUB6 line. Microtubules are enriched on neck sides while comparatively scarce on the lobe sides of the anticlinal walls. These microtubules seem to be continuous between periclinal and anticlinal walls. Related to Fig. 6B, C and S7D.

**Movie S4**. 3D reconstruction of confocal z-stack demonstrating microtubules (orange, GFP-MAP4) and cell wall (green, propidium iodide) in pavement cells of *Arabidopsis* cotyledon. Marking the cell borders with the green channel (appears and disappears during the movie) reveals that microtubules are enriched on the neck sides. Related to Figs. 6A, 7C-F and S7E, F.

**Movie S5**. 3D reconstruction of confocal z-stack demonstrating microtubules in pavement cells of *Arabidopsis* cotyledon from GFP-TUB6 line. Arrows show microtubules extending between the periclinal and anticlinal walls at neck locations of tiny bends formed in the cell wall. Related to Figs. 6B, C and S7D.

## Supplemental Information

Available in the online version.

## Notes

#### Summary of Updates

Addition of Figures 1E and F. Earlier Figure 2 is moved to Supplemental Information. Minor edits throughout.

