## Supplemental Information for "Mechanical stress initiates and sustains the morphogenesis of wavy leaf epidermal cells"

### Supplemental Notes

#### The mechanics of the anticlinal wall alone is not sufficient to describe the development of interlocking lobes

Anticlinal walls of epidermal pavement cells form the cell borders and their wavy outlines observed under the microscope are the most conspicuous aspect of the interlocking morphogenesis of pavement cells. Since the waviness of these walls is the feature that defines the 2D representation of cell shapes, many studies have based their focus mainly or predominantly on the anticlinal walls. While such simplification can be acceptable in some cases, depending on the nature of problem being investigated, it can also result in misrepresentation of the cell mechanics and shape morphogenesis. Here we test two concepts based on isolated anticlinal wall models that would have the potential to be morphogenetic agents: differential turgor and differential mechanical properties in the adjacent layers of the anticlinal cell wall. In addition to these analyses, we also performed a separate but related analysis which revealed that a model based on only anticlinal border walls forming undulations based on "tissue tension" as proposed by Majda et al. (2017) results in conflicting outcomes (Bidhendi and Geitmann, 2019).

###### Supplemental Note 1

#### Generating cellular protrusions based on anticlinal wall modification requires differential turgor pressure

To determine which mechanical conditions would be required to generate a bend in the anticlinal wall resulting in the protrusion of one cell into the neighboring cell, we isolated the problem and developed a finite element model of a section of anticlinal wall between two adjacent cells (Fig. S1A). For this isolated approach, periclinal walls were neglected. The anticlinal wall was constructed to consist of two layers (the primary walls of the two adjacent cells) that are glued together (by the middle lamella, a pectin-rich, thin layer that connects cells in a plant tissue).

The first hypothesis we tested assumes that the curvature in the anticlinal wall is generated because of different growth rates of the two layers, similar to a temperature controlled actuator consisting of two layers with different thermal expansion properties or in curling of thin bilayers (Pezzulla et al., 2016) (Fig. S1A). Relative displacement between the layers is not allowed to mimic the role of the middle lamella. One layer is set to expand at a higher rate by assigning a higher coefficient of thermal expansion—our modeling equivalent of the expansion of cell wall material through the addition of softer material. The simulations show that the layer at the convex (neck) side of the forming bend is the one with the higher rate of expansion (Fig. S1B). However, this contradicts our understanding of the biology of pavement cells. The neck side is thought to be stiffened by microfibrils (see Figs. 1D, 6 and Movies S1 and S2), whereas a putative accumulation of actin arrays at the concave side of undulations is suggested to deliver new matrix material by exocytosis predominantly on the lobe side (Panteris and Galatis, 2005). The finite element model based on asymmetric anticlinal wall expansion, therefore, seems unable to provide a description of wave formation that is consistent with the known biological processes.

The second hypothesis we tested is not based on modulation of wall growth behavior, but on that of the acting forces. We examined whether differential turgor in neighboring cells can generate a bend in the anticlinal wall. For modeling purposes, the cell geometry is hexagonal (Figs. S1C). As above, non-slipping non-separable contact is considered between the anticlinal wall layers in contact. This is to reflect the adhesion between the cell wall layers at the middle lamella. Turgor pressure is exerted on the internal surfaces of both cells, and the model allows for independent adjustment of the turgor pressure in each cell.

At equal turgor pressures in the two neighboring cells, the overall shape and the relative position of the midline of the anticlinal wall in contact between two cells does not change as the forces acting on the two sides cancel each other out. If differential turgor pressures are applied, the cell with the higher pressure curves into the cell with lower pressure (Fig. S1D). A more spatially confined deformation similar to a lobe protrusion can be achieved if the anticlinal wall layers are made softer on a limited length, combined with the application of differential pressure (Figs. S1E and F). To produce a local protrusion, this locally reduced stiffness must be applied to both wall layers in the same region (although not necessarily at the same magnitude). If only one wall layer is softened, the deformation resulting from the differential pressure application causes an overall bend of the entire wall section rather than a spatially confined protrusion.

**Supplemental Note 2**

#### Mechanical changes confined to anticlinal walls are not sufficient for cell growth

To investigate whether the predictions made by the isolated anticlinal wall model remain valid in the presence of the periclinal walls, these are added to the finite element model (Figs. S1G and H). The aspect ratios for cell wall dimensions are adopted from typical values obtained from pavement cells in *Arabidopsis* cotyledons at early stages of development (e.g., Fig. S5G). Periclinal wall thickness is set to be identical to that of the anticlinal walls, and the material is configured to be isotropic with uniform stiffness. The pressure in the right cell is set 10:1 compared to the cell on the left. The simulations show that upon application of the turgor pressure, the cells swell out of the plane, the free anticlinal walls displace inward, reducing the in-plane cell surface (Fig. S1H). The anticlinal wall shared with the neighboring cell displaces toward the cell with higher pressure (Figs. S1I and S1K). Softening of the whole anticlinal wall in the contact region between the cells results mainly in an upward stretch of this wall, a displacement toward the cell with a higher pressure with a bulge in the mid-depth of the anticlinal walls toward the cell with lower pressure. However, the superficial borders of the anticlinal and periclinal walls remain relatively straight (Fig. S1J). A high ratio of turgor pressures (100:1) was used to produce a discernible behavioral trend. Even at this unphysiologically high pressure differential, the protrusion remains isolated to the anticlinal wall and forms no visible wave at the superficial borders of the cells. It should be noted that in the biological system, even relatively small turgor differentials between adjacent cells may be difficult to achieve or maintain due to the existence of intercellular channels, plasmodesmata, which allow for a certain degree of fluid exchange. Similar results are obtained when the softening is confined to an isolated region of the contact anticlinal wall. Therefore, unlike the isolated anticlinal wall model, if cell walls are mechanically isotropic, pressurization of a cell does not generate behavior expected from a differentiating epidermal cell. It neither increases the area of the cell in plane nor does it generate anticlinal curvatures into the adjacent cells. These results can be explained considering the respective surface areas of the periclinal and anticlinal walls. When pressurized, the geometry of any body tends to swell in a manner that transforms its shape closer to spherical. In the case of the tabular pavement cells of the *Arabidopsis* cotyledon, this occurs since the combined inner and outer periclinal walls possess a higher surface area compared to the combined anticlinal walls of the same cell (see for instance Fig. S5G). As a consequence, the force of the turgor pressure acting upon the periclinal walls ($F = P \times A$, where $P$ is the pressure and $A$ is the surface area on which the pressure is exerted) is greater than that acting on the combined anticlinal walls resulting in a net out-of-plane deformation of the cell. This is not to consider that, regardless of the difference in surface area, the effect of turgor pressure on anticlinal walls can be cancelled out when the pressures inside neighboring cells are equal. The out-of-plane swelling of the periclinal walls leads to a contraction of the in-plane dimensions of the cell. These simulations therefore, also suggest that isotropic softening of the anticlinal walls does not cause epidermal plant cells to grow in plane.

### Supplemental Modeling Procedures

##### Supplemental Note 3

#### Material and model inputs

For all simulations, neo-Hookean hyperelastic material model was used to define the cell wall material behavior. We constructed our finite element models with normalized (dimensionless) inputs. Dimensionless values facilitate assessing the model behavior without the need to operate on very small numbers that arise due to microscale dimensions of the geometries. This also reduces the possible round-off errors that may arise due to operations on very small numbers during numerical calculations.
Two material constants are needed to define a neo-Hookean material model in Abaqus (Abaqus, 2014):
$C_{10} = \frac{\mu_0}{2}$ and $D_1 = \frac{2}{K_0}$
$\mu_0$ and $K_0$ correspond to initial shear and bulk moduli, respectively. The material constants were normalized by $\mu_0$. Therefore, for a region with default stiffness, a $C_{10} = 0.5$ could be used. Reduced or increased stiffness were similarly input by inputting values less or more than $C_{10} = 0.5$, respectively. Values of $C_{10}$ used to test the behavior of the models varied between $5 \times 10^{-2}$ and $50$. The material in all models was considered incompressible. For such a case, $D_1 = 0$. It should be noted, however, that compressibility would not affect the behavior of the model if it was input otherwise. The turgor was applied as a distributed pressure on internal surfaces of the cell walls. The turgor pressure input was also normalized by $\mu_0$ (same units). The range of dimensionless values used for the turgor pressure varied between $1 \times 10^{-5}$ to $1 \times 10^{-2}$. The dimensions of the cell geometries were also normalized with respect to one of the dimensions. The base value for cell wall thickness was approximated as 700 nm from images and was rounded up to 1 μm. The height of the anticlinal wall and its length were also approximated to 10 and 100 μm, respectively. For solid models, the thickness, height and length for the solid model with only the anticlinal wall included were $0.01$, $0.1$, and $1$, respectively. For shell models with a piece of the cell wall containing multiple lobes and necks or for whole cell models, these dimensions were 0.1, 1 and 10, maintaining the same aspect ratio.

##### Supplemental Note 4

#### Finite element implementation

For the model including only the anticlinal wall, continuum quadratic three-dimensional large strain elements with reduced integration (C3D20RH) were used. The anticlinal walls of two adjacent cells were tied together in all degrees of freedom. For beam models, linear two-node beam elements were used. In other models, the cell wall was considered as thin shell as the thickness of the cell wall compared to other cell dimensions is negligible. These models were discretized with four-node first order reduced integration shell elements (S4R). In multi-cell models, the geometries of individual cells were merged at their anticlinal walls. Boundary conditions were applied to allow free deformation of wall segments under turgor pressure while preventing movement of the whole body in 3D space.

To compare deformations in ab initio or cum tempore stiffening models, the secondary onset of stiffening in a segment of cell walls in the cum tempore model was applied indirectly through exploiting a temperature-dependent stiffness scenario. As temperature can be defined to change in time, the stiffness could be made to vary in a time-dependent manner. Quantitative comparisons between ab initio stiffening and *cum tempore* stiffening were performed by monitoring the displacement of four fiducial points on the 2D shell model (Fig. S2C). The results indicate that in both cases, similar displacements for anticlinal and periclinal walls result from the application of turgor (Figs. S2E and F).

**Supplemental Note 5**

#### Construction of the buckling model

Proof-of-concept buckling models were developed to demonstrate that the cell walls, including both the anticlinal and periclinal walls, can buckle resulting in wavy cell contours. For this, the cell was modeled as a hollow rectangular box. Shell behavior was considered for the cell walls. Linear elastic material was used for linear buckling analysis. We observed that with and without a static preloading step, the structure can buckle under internal pressure and positive eigenvalues for the buckling analysis were found. The eigenvalues of a buckling analysis depend greatly on the geometry, dimensions and material inputs as well as the boundary conditions. The image provided in the manuscript was from a model with dimensions of 100, 10 and 1 μm for length, height and the shell wall thickness, respectively. The material was considered as linear elastic with a Young's modulus of 1 MPa and Poisson's ratio of 0.3. Boundary conditions were applied to prevent the in or out of plane displacement of the inner (lower) periclinal walls (as attached to mesophyll cells). However, further simulations showed that the outcome in terms of feasibility of buckling is not dependent on this particular boundary condition. Turgor load was applied to inner faces of outer periclinal walls. For these inputs, critical buckling load for turgor pressure was found to be as low as 2.8 kPa which is well below the reported range for turgor pressure in plant cells.

**Supplemental Note 6**

#### Construction of the stress-stiffening feedback loop

To implement a feedback loop, a Python script was developed to read and write in the finite element model (Fig. S6). The code is available upon request. After each iteration, the code extracts the deformed geometry from the Abaqus database and reads the stresses for each element. If a specific element has a stress higher than a threshold and does not belong to a list of stiffening-inhibition zone (in model accounting for inhibition of stiffening), the new value of stiffness for that element in terms of $C_{10}$ is updated according to $C_{10_{new}} = C_{10_{old}} + tanh(C_{10_{old}} \times S_{rel})$; where $S_{rel}$ is the von Mises stress of the corresponding element relative to the threshold stress. The threshold stress was calculated in each iteration as average stress of all elements. Otherwise, $C_{10_{new}} = C_{10_{old}}$ was assigned for the element with either a stress below the threshold stress or located in a stiffening-exclusion region. After assigning new stiffness values, the script runs the model keeping all other model parameters such as the turgor pressure and boundary conditions as the previous iteration.

### Supplemental Experimental Procedures

**Supplemental Note 7**

#### Polysaccharide staining

To visualize pectin status, seedlings were stained with 0.5 mg/mL aqueous solution of propidium iodide in double distilled water (dd$H_2O$). A drop of propidium iodide was placed on each seedling. The dye was removed after 10-20 minutes with a Kimwipe tissue paper and washed gently with double distilled water (dd$H_2O$) at least three times before mounting in water for visualization. For COS$^{488}$ staining, the stock was diluted 1:500 in MES buffer (25 mM, pH 5.7). The seedlings were incubated with COS$^{488}$ for 5-15 minutes and washed with MES for 3-5 times for at least 30 seconds. The samples were then mounted in MES before visualization. For visualizing cellulose, calcofluor white and PFS were used. Calcofluor was used at a concentration of 2 mg/mL in dd$H_2O$. Seedlings were placed in Eppendorf tubes containing calcofluor white and transferred to a vacuum of 20 in Hg (Pelco BioWave 34700) at room temperature. After 45 minutes the tubes were transferred on a rotator in a dark room for an additional 45 minutes. The specimens were then washed gently for 3-5 times with $ddH_2O$ before being mounted in $ddH_2O$ for observation. PFS staining was carried out with a 14 mg/mL solution of PFS in PBS buffer ($Na_2HPO_4$ 3.2 mM, $KH_2PO_4$ 0.5 mM, NaCl 135 mM, KCl 1.3 mM, pH 7.3). The staining of specimens was performed as described above for an incubation time of 30-60 minutes before washing and mounting in PBS for observation.

### Experimental measurements and statistical analysis

##### Supplemental Note 8

#### Pectin signal intensity

$COS^{488}$, shown to be highly specific to weakly esterified pectin (Mravec et al., 2014), was used to stain pectin. Using ImageJ, maximum signal intensity projections were obtained from confocal z-stacks. Like measurements carried out for comparison of GFP intensity from microtubules between necks and lobes, circles were placed on two sides of a wave on necks and lobes to compare signal intensity from $COS^{488}$ staining (Fig. S7H). The area of the circle was kept constant between all measurements (5 $\mu m^2$). Images of 5 cotyledons from 5 different seedlings were chosen and from each cotyledon 1-2 pavement cells were analyzed for $COS^{488}$ signal making in total 55 pairs of neck and lobe measurements. Paired t-test between data acquired from lobes and necks showed a significant difference as necks appeared consistently brighter than lobes ($p<0.0001$).

Analysis of propidium iodide signal intensity for pectin staining was carried out as described above for $COS^{488}$ staining. 5 cotyledons from different staining experiments were used and for each, between 3 to 5 pavement cells were chosen for analysis making up to 19 cells in total and a total of 101 lobe/neck pairs that were analyzed. Signal intensities obtained for necks were in majority of cases higher than lobes and the differences were significant as determined in paired t-tests ($p<0.0001$).

##### Supplemental Note 9

#### Cellulose signal intensity

Calcofluor white and Pontamine Fast Scarlet 4B (PFS) were used to label cellulose, as described in the polysaccharide staining section. Fibrillar bundles of cellulose were prominent fanning out on the indentation (neck) sides (Figs. 6D, E, F, S5D-F. Movies S1 and S2). Extension of cellulose enrichment at locations of necks could also be observed down the anticlinal walls (Fig. S5F1 and F2, Movies S1 and S2). To ensure that this observation is consistent, signal intensities in maximum z projections were compared between necks and lobes of periclinal walls in PFS stained samples since this dye is suggested to be highly specific in binding cellulose (Anderson et al., 2010; Liesche et al., 2013). Similar to the measurements described above for pectin, this was carried out by placing circular regions of interest in which mean signal intensities were read and compared. 6 cells from 3 seedlings were selected from which a total of 88 lobe/neck pairs were analyzed. A paired t-test showed a significance difference between signal intensities of necks and lobes ($p<0.0001$). This pattern was similar to the observed microtubule enrichment in the necks under periclinal walls.

**Supplemental Note 10**

#### Microtubule localization

##### Cortical microtubules at the periclinal wall are more abundant in necks than lobes

Periclinal microtubule density was compared between necks and lobes of pavement cells. This was carried out by reading the mean signal intensity in a circle with a consistent area (8.5 μm$^2$) placed on two sides of the cell border of a wave (Fig. S7D). z-stacks of pavement cells of GFP-TUB6 line were acquired for cotyledons of seedlings between 2-5 days after germination. The depth of Z-scanning was typically between 10 to 20 μm and was adjusted for each z-stack by determining the first and last slices of the stack to ensure that the outer periclinal walls, the entire height of anticlinal walls, as well as at least some portion of the inner periclinal walls were captured to allow for a 3D view of the cell structures. This approach allowed us to later dissect the z-stacks and study the outer periclinal walls and the anticlinal walls separately. For each micrograph used in analysis, a slide was determined from the z-stack that contained all cell borders (top of the anticlinal walls) with little to no periclinal wall signal showing. These particular sections were pseudocolored orange/red to mark the cell borders (e.g., Fig. S7D). To collapse the periclinal wall information into a single image, maximum projection of the fluorescence signals from the first optical section down to the specific section chosen as beginning of the anticlinal wall was made and pseudocolored yellow. The borders colored red/orange and the max projection data from the periclinal wall were merged into a single micrograph as depicted in Fig. S7D. These micrographs were used to compare signal intensity on the opposing sides of borders in circular regions of interests on necks and lobes (Fig. S7D). 6 pavement cells were analyzed from 6 different cotyledons. 84 lobe/neck pairs were analyzed. In the majority of cases, the mean signal intensity was considerably higher on the neck side and t-test was carried out on all the pairs with ($p<0.0001$).

##### Cortical microtubules at the anticlinal wall are more abundant at the neck sides, extending from the periclinal microtubule enriched zones

As described in the previous section, we observed that cortical microtubules under periclinal walls are consistently enriched at the neck regions of undulations. We hypothesized that this pattern of microtubule enrichment persists in neck regions of the anticlinal walls as well. To verify whether a significant difference exists between the abundance of microtubule at the neck and lobe sides of the anticlinal walls, we counted the cortical anticlinal microtubules on each side. This was carried out using 3D reconstructions of z-stacks of GFP-MAP4 line cotyledons between 2-5 days after germination. Specifically, using overlapping double channel 3D reconstructions marking microtubules (GFP-MAP4) and cell wall polysaccharides (propidium iodide) allowed us to ascertain whether a given microtubule array lined the neck or the lobe sides (Movie S4, Fig. S7F). We analyzed 35 lobe/neck pairs from 9 different pavement cells randomly selected from z-stacks of 3 cotyledons of the GFP-MAP4 line. Segments with arc length of 6 μm were selected choosing points on the cell borders on maximum projection micrographs using Fiji plugin Kappa (developed by the Brouhard laboratory, brouhardlab.mcgill.ca) (segments of the anticlinal walls, Fig. 7SE). A B-spline was fitted to the selected points. Kappa provides the length of the chosen arc and its average curvature. Curvature is $\frac{1}{R}$, where $R$ is the radius of a circle fitted to the arc. Analysis was carried out on a range of wall segment curvatures starting from relatively straight borders to rather pronounced lobes. Paired t-test showed that the microtubule population is consistently denser on the neck sides ($p<0.0001$). Further, we observed a moderate correlation ($r=0.5$), when relative difference in microtubule number between necks and lobes in percentage ($\frac{N_{neck}-N_{lobe}}{N_{neck}} \times 100$) was plotted against segment curvatures (Fig. S7G). Pronounced bends showed a higher difference in microtubule numbers on the two opposing sides of the anticlinal wall. Obviously, the progression of a lobe does not depend on line curvature alone and further studies could take into account the depth (base to tip length) of the lobes in addition to their curvature for a more thorough analysis of the correlation between shape and stage of wall curvature and microtubule polarization. However, this was beyond the scope of the present paper. The scoring of microtubule location with respect to neck or lobe side of the anticlinal wall was based on the position relative to the anticlinal wall, corroborated by the fact that the arrays continued from the anticlinal wall region to radiate under the periclinal wall of the respective cell. This was particularly obvious in neck regions (Movie S3).

**Supplemental Note 11**

#### Measurement of dimensions of growing cells from time-lapse data

In order to determine the effect of CGA application on pavement cell shape, cell circularity, area, and perimeter were measured at 2 time points between 2 and 4 days after germination. Cell perimeters were traced manually in ImageJ, on the z-stack projection of the cells. To ensure the highest possible precision, we eschewed the use of automated algorithms but instead counted lobe numbers manually (the 'gold standard' against which most automated algorithms are compared). 50 to 70 cells from 10-12 seedlings were used to produce each data point. A paired t-test showed a significant difference in the lobe number between the control and treated cells ($p<0.001$). Details of CGA treatment can be found under Experimental Procedures of the main manuscript.

#### Supplemental Figures

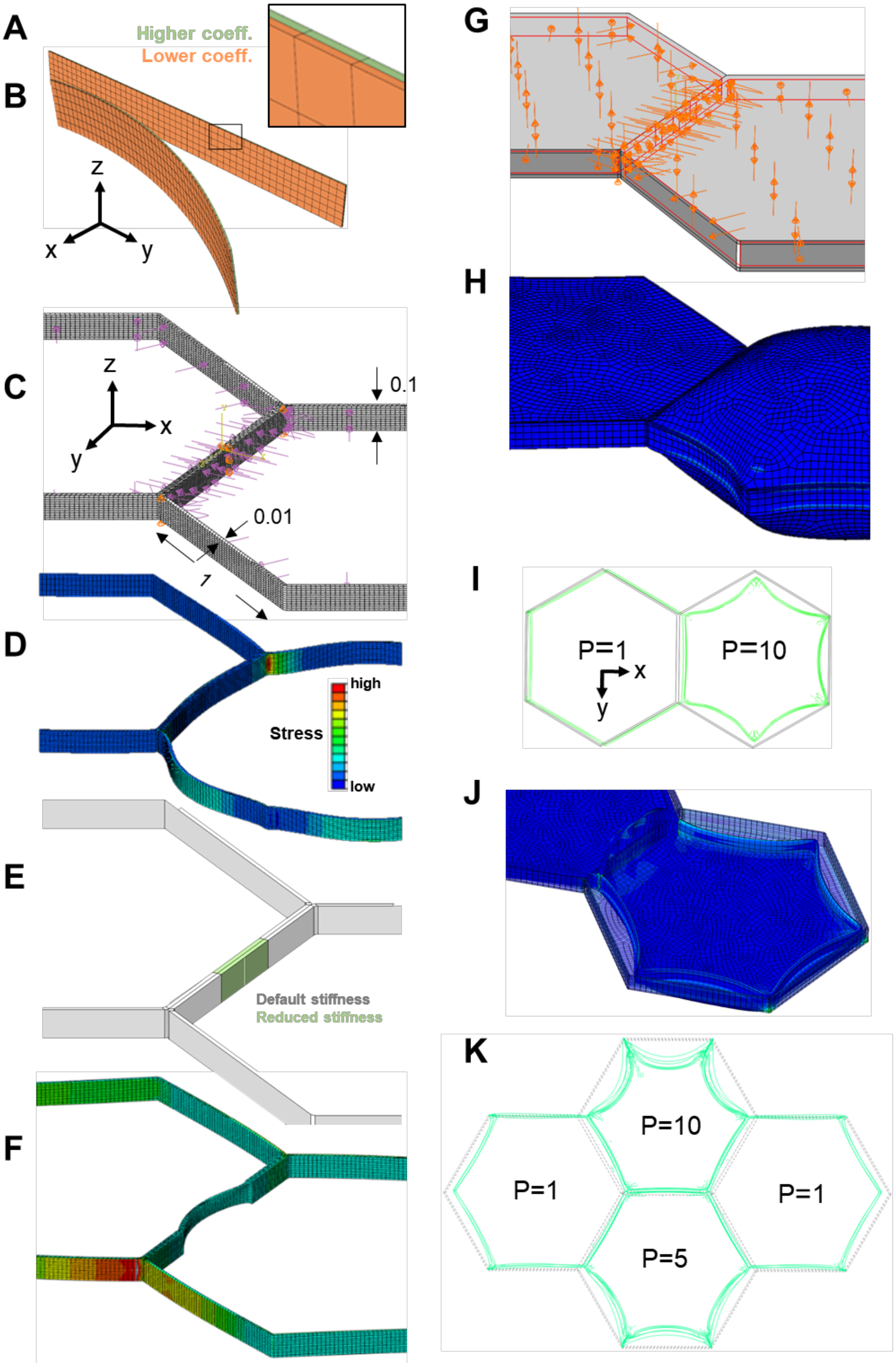

**Fig. S1**. Related to Figs. 2 and 3, **finite element models designed to investigate the effect of anticlinal wall stiffness modulation and turgor pressure differential on cell shape**. **A)** Representation of anticlinal wall separating adjacent cells by two material layers glued together at their interface. One layer (green) is given a higher coefficient of thermal expansion to simulate higher growth rate; the right end of the wall may move in space. **B)** Differential expansion of the layers results in the formation of a curvature with the faster-expanding layer forming a convex curve. **C)** Relative dimensions in the model focusing on only the modulation of anticlinal wall between adjacent cells and the turgor pressure. Pressure loads are applied on internal surfaces of the walls. **D)** In the presence of differential pressures and uniform anticlinal wall stiffness the anticlinal wall forms a second order curve into the cell with lower pressure. Heatmap represents stress. **E)** A spatially confined region (green) in the anticlinal wall is assigned a softer value than the default stiffness (gray). **F)** A turgor differential results in the formation of a local protrusion into the cell with lower turgor. **G)** Periclinal walls are added to the model. **H)** With isotropic material properties for all cell walls, the periclinal cell wall of the cell with the higher turgor pressure forms a more pronounced swelling in z direction pulling the anticlinal walls inwards. **I)** Displacement of cell borders shown from above for two adjacent cells with isotropic material properties and identical stiffness in anticlinal and periclinal walls with different turgor pressures. A pressure difference with ratio of 1:10 was implemented. Upon application of turgor pressure, due to out-of-plane swelling of cells, in-plane cell dimensions contract with the anticlinal wall shared with the neighboring cell displacing and forming a slight curvature toward the cell with a higher turgor pressure. **J)** Inside view: The shared anticlinal walls between two cells is softened compared to the other walls in the model. The outer periclinal walls are removed from the view to show the status of the anticlinal wall. The softened anticlinal wall forms a bulge toward the cell with a lower pressure but the top and bottom edges with the periclinal walls remain straight. **K)** Model of four cells sharing two tricellular junctions, with different relative values of turgor pressure (P)**.** Turgor induced deformation of the anticlinal walls viewed from the top. The construct of this model is the same as the model in (**I**) but was intended to study the behavior of anticlinal walls embedded between several cells and not located at the free edges.

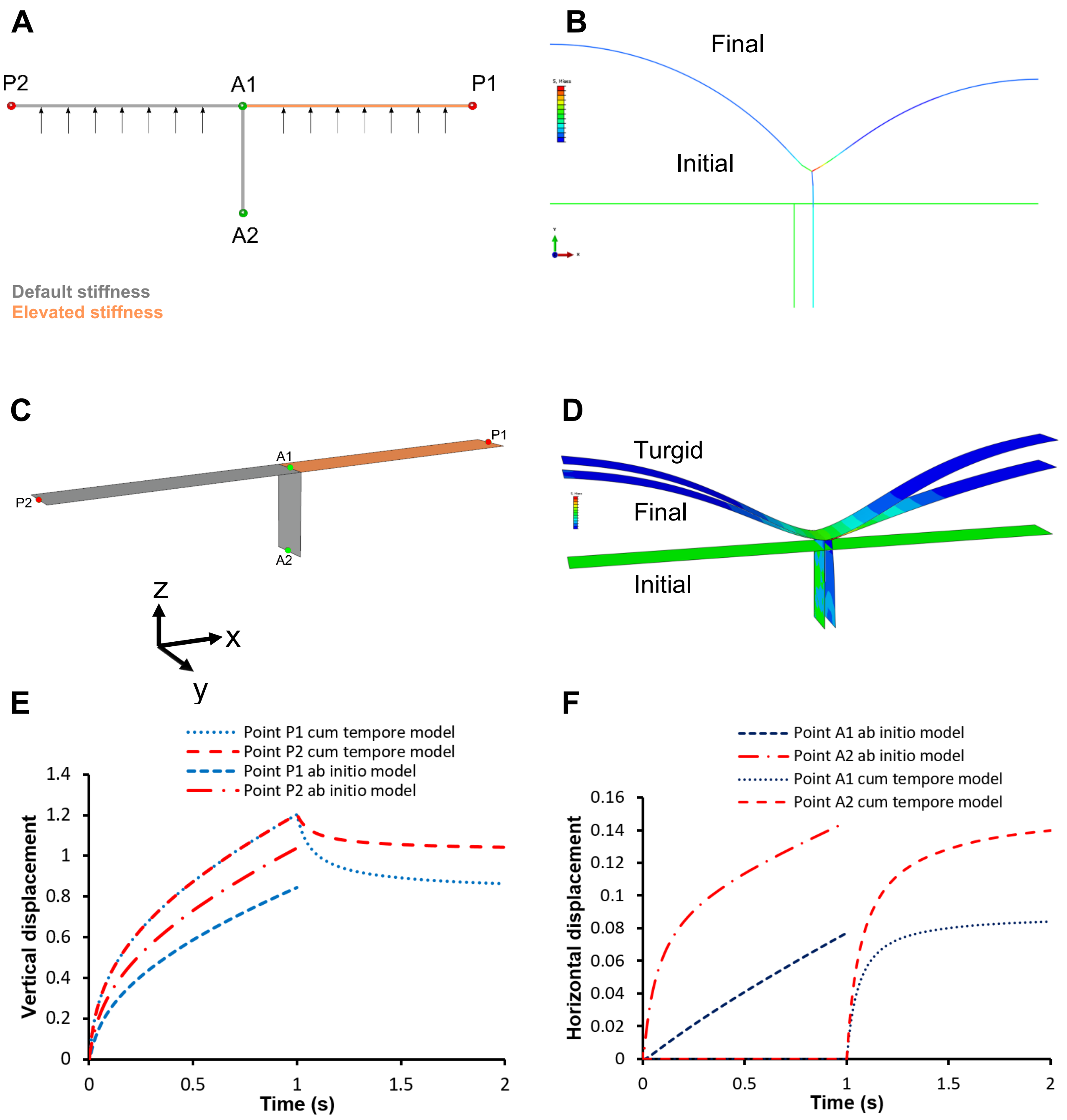


**Fig. S2.** Related to Fig. 3, **beam and shell models of wall segments. A)** 2D beam model of the wall segment demonstrated in Fig. 3A. Turgor pressure is applied on internal edges of the beam elements. At the anticlinal wall the effect of equal turgor on both sides cancels each other out, but periclinal wall segments bulge out of the plane of the epidermis. Four points of interest on periclinal and anticlinal walls are identified for recording the resulting displacements. **B)** Initial and deformed shapes of the 2D model showing displacement of the anticlinal wall segment toward the stiffer (neck) side. Finite element models of wall segments implementing *cum tempore* stiffening. The onset of stiffness augmentation is applied when the walls are already under tension due to turgor pressure. **C)** Shell model for *cum tempore* stiffening. Pressure is applied to the inner side of the periclinal walls and the right periclinal wall segment is stiffened after the full application of pressure. **D)** Displacement for the shell model resulting from repeated pressure application with *cum tempore* stiffening. The anticlinal wall is displaced toward the side with the stiffer periclinal wall. **E** and **F)** Comparison of the displacement of fiducial markers on shell model for *ab initio* and *cum tempore* models.

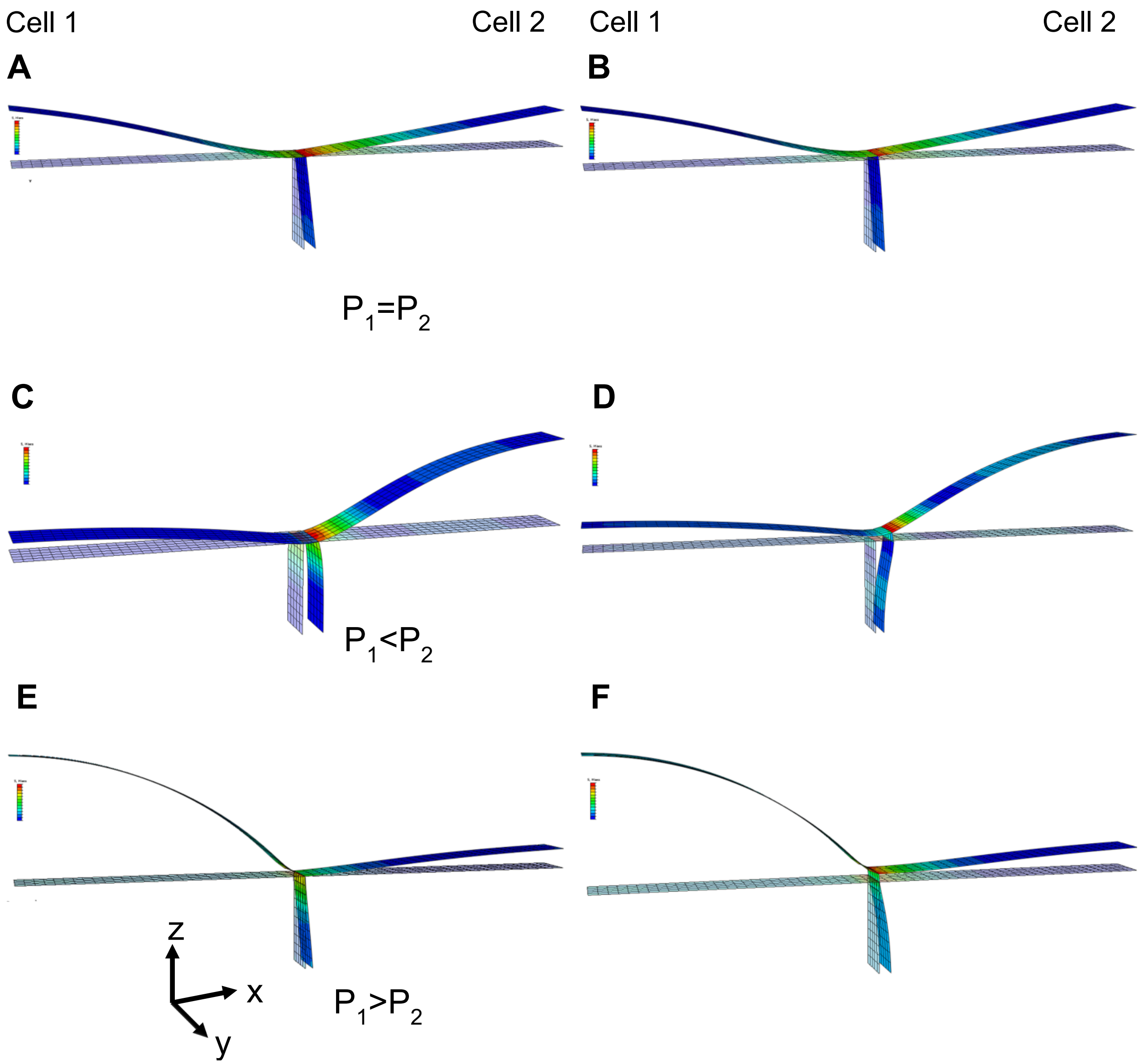


**Fig. S3.** Related to Fig. 3, **shell model of segments of periclinal and anticlinal walls with stiffened periclinal wall** in cell 2 (normalized, $C_{10}$=2). Turgor pressure is either identical in both cells (**A, B**), higher in cell 2 (**C, D**) or in cell 1 (**E, F**). Anticlinal wall is either stiffened ($C_{10}$=2, in **A, C, E**) or has default stiffness ($C_{10}$=1 in **B, D, F**). In all cases is the anticlinal wall displaced towards cell 2.

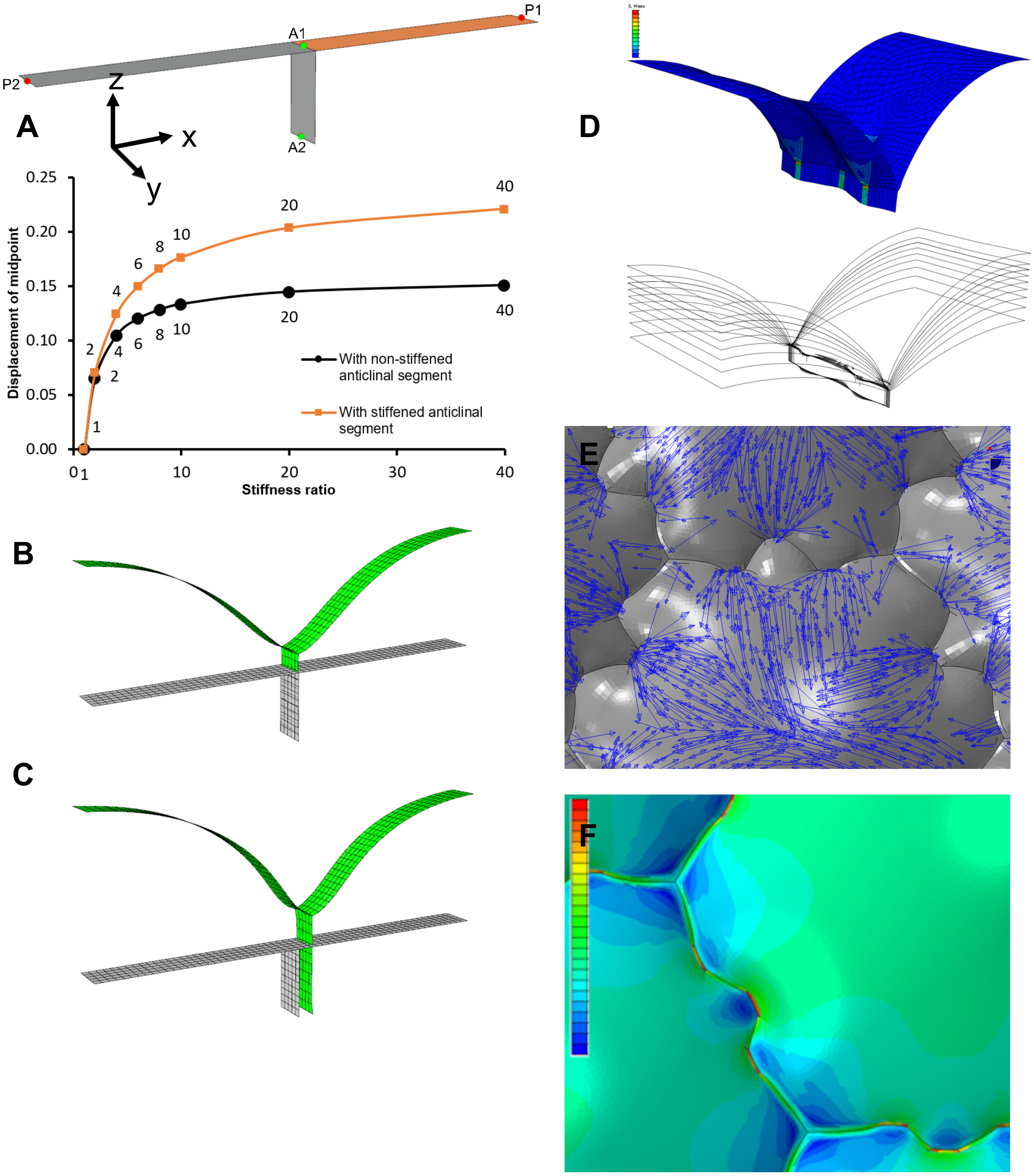


**Fig. S4.** Related to Figs. 3 and 4, **partial and whole cell finite element models of cell wall deformation**. **A)** Horizontal displacement of Point A2 against the stiffness ratio between the periclinal wall segments with default and increased stiffness, with stiffened and non-stiffened anticlinal wall segment. **B)** Displacement of the anticlinal wall after three iterations of load application with stiffness ratio between the right and left periclinal segments equal to 1.02. **C)** Ditto with stiffness ratio equal to 2. **D)** Evolution of undulations for the model with alternate placement of stiffenings on periclinal and anticlinal walls by iterating load application with relieving the wall stress after each iteration. **E)** Stress field reveals stress concentration at necks and cell corners. **F)** Geometry with undulations at cell borders used as an input geometry with isotropic material properties and same stiffness value in all regions of the model. The stresses are higher on the convex side of undulations.

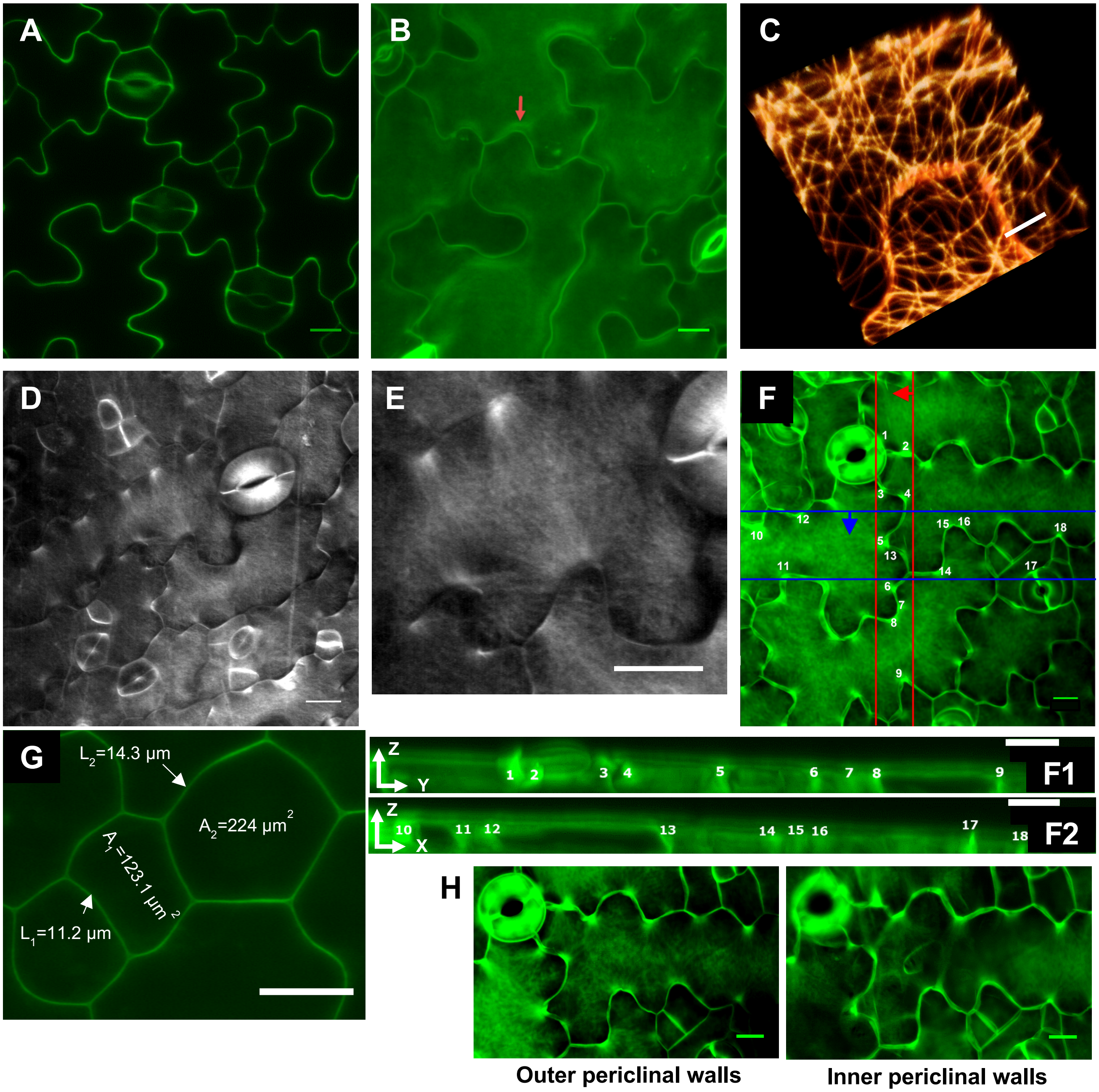


**Fig. S5.** Related to Figs. 5-7, **fluorescence micrographs of *Arabidopsis thaliana* cotyledons at 3 to 5 days after germination**. **A)** Single optical section through middle of the epidermal layer thickness, showing the borders of cells stained with propidium iodide, reveals varying signal intensity along the length of anticlinal walls. Tri-cellular cell junctions typically display high signal. **B)** Maximum projection of z-stack reveals propidium iodide signal intensity in periclinal walls to be higher at neck locations. **C)** Microtubules labeled with GFP-MAP4 show more abundant bundling on the neck side, although the bundles can also occasionally be found extending to the tips of lobes. The image represents an oblique view of a confocal z-stack. **D)** and **E)** Pontamine fast scarlet 4B reveals localization of cellulose bundles in periclinal walls at neck sides of undulations. **E** shows magnified region from **D** as indicated. **F)** xy maximum projection of z-stack stained with calcofluor white**: F1)** yz and **F2)** xz projections of the cell walls between the lines marked on figure **F**. **G) I)** Typical dimensions of epidermal cells in the *Arabidopsis* cotyledon. $L_1$ and $L_2$ are the lengths of anticlinal walls with corresponding surface areas of 58.2 and 74.4 μm$^2$, respectively. Their surface was measured based on the depth of the anticlinal wall measured to be 5.2 μm (obtained from the corresponding z-stack). Areas $A_1$ and $A_2$ correspond to the surfaces of outer periclinal walls of the two cells. **H)** Comparison of cellulose orientation in outer and inner periclinal walls of the same cell using calcofluor white. Images are produced by maximum projection of upper and lower halves of z-stack data, separately. Scale bars = 5 μm (**C**) and 10 μm (**A**, **B**, **D**, **E**, **F, G** and **H**).

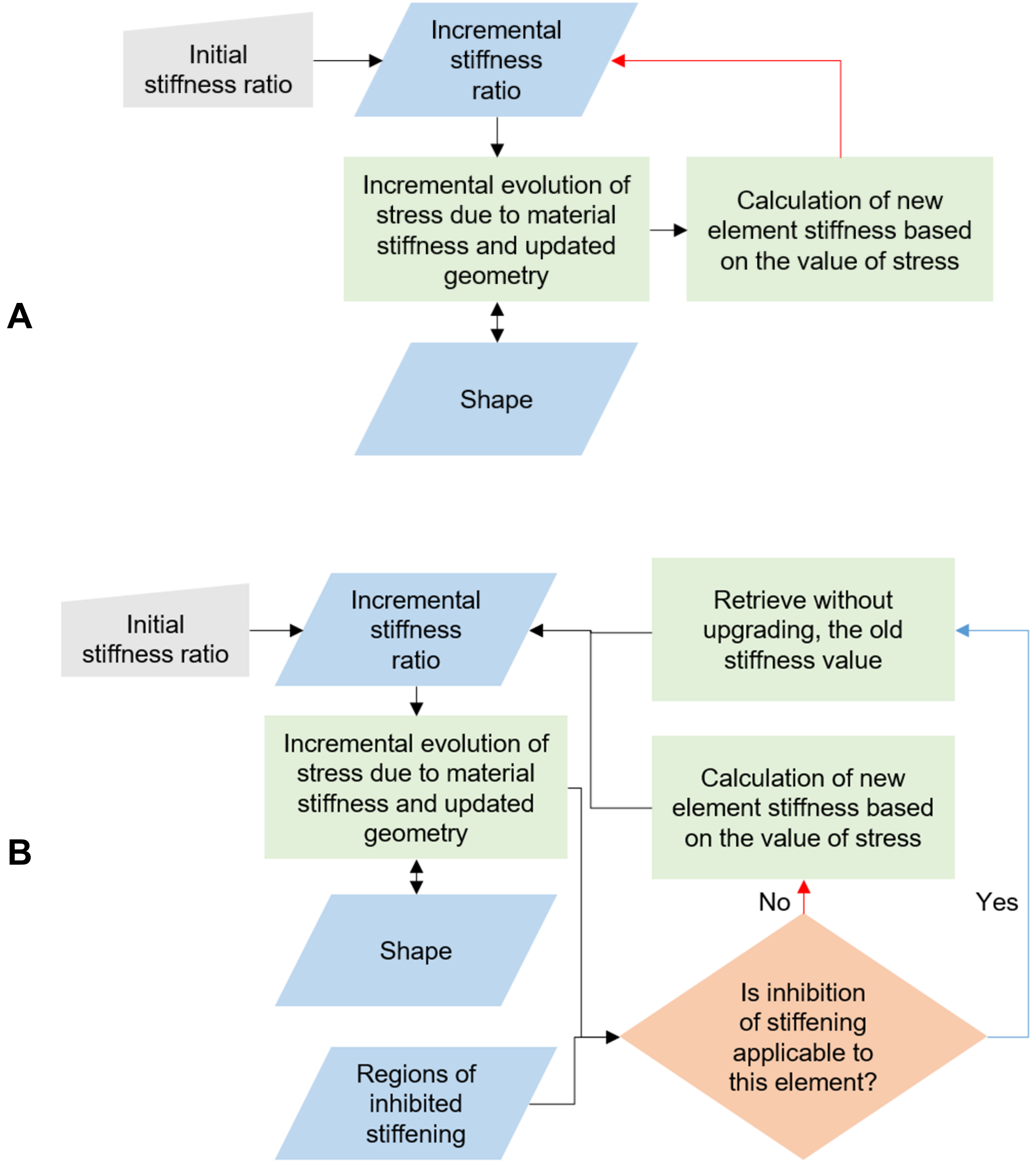


**Fig. S6.** Related to Fig. 4, **steps in the positive mechanical feedback model linking the stresses, wall deformation and local stiffness**. **A)** Without inhibition of stress-induced stiffening in regions alternating with incipient necks. **B)** Feedback model implementing inhibition of stiffening in regions alternating with incipient necks.

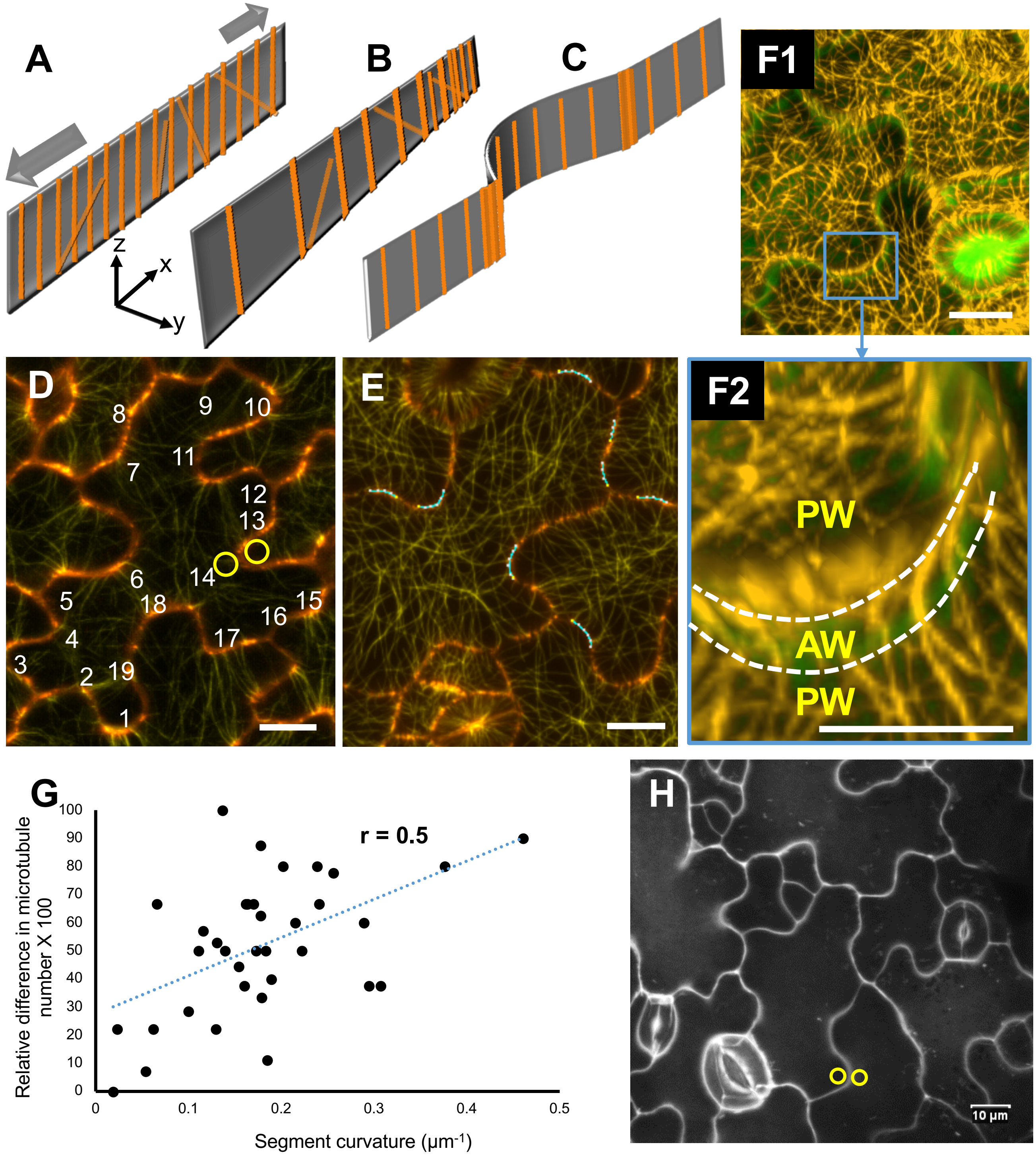
A
B
C
F1
z
x
y
D
8
9
10
7
11
12
13
6
14
5
18
15
16
4
17
3
2
19
1
E
F2
PW
AW
PW
G
r = 0.5
Relative difference in microtubule number X 100
Segment curvature (µm-1)
100
90
80
70
60
50
40
30
20
10
0
0
0.1
0.2
0.3
0.4
0.5
H
10 µm

**Fig. S7.** Related to Figs. 5-7, **spatial distribution of microtubules and pectin A**) Preferential alignment of microtubules along undulations on the anticlinal walls suggesting that subsequent deposition of cellulose microfibrils renders the wall mechanically transversely isotropic. **B**) As a result of microtubule and cellulose orientation, the anticlinal wall can expand similarly to an accordion in plane, while out of plane (z direction) expansion is restricted. **C**) Progression of a lobe is accompanied by marked accumulation of microtubules and presumably increased deposition of cellulose at its two base-points (or necks of the adjacent cell). **D**) Signal intensity for cortical microtubules underlying the periclinal wall was measured at necks and lobes using maximum intensity projections of z-stacks acquired collecting the GFP-TUB6 signal. Cell borders were pseudocolored red/orange by superimposing an optical section from the middle of the stack using the propidium iodide signal. **E**) Segments of an undulation were chosen in 2D maximum projection micrographs, obtained as in (D) for GFP-MAP4, to mark regions for microtubule count in 3D. Kappa Fiji plugin (Brouhard laboratory, 2016-17) was used to mark and measure the curvature of line segments. **F**) Oblique view of **F1)** 3D reconstruction of two-channel data in Imaris. Green pseudocolor shows the cell wall (propidium iodide) and the yellow channel represents the microtubules based on GFP-MAP4 signal. **F2)** Zoom-in oblique view of the region of interest from (F1), with slightly more tilt. The dotted line is placed to indicate the cell border at which anticlinal (AW**)** and periclinal (PW) walls connect. 3D reconstruction in Imaris was used to count microtubules adjacent to the anticlinal walls. **G)** Relative difference in microtubule number presented in percentage plotted against the curvature of the segment used to define the region of interest for counting (E). A moderate correlation (r=0.5) was found, with the difference in microtubule abundance between lobes and necks increasing with wall bend curvature. **H)** Maximum projection of z-stacks from $COS^{488}$ staining. Circular regions of interests were placed on two sides of border waves to measure mean signal intensity for necks and lobes on the periclinal walls. Scale bars = 10 μm.

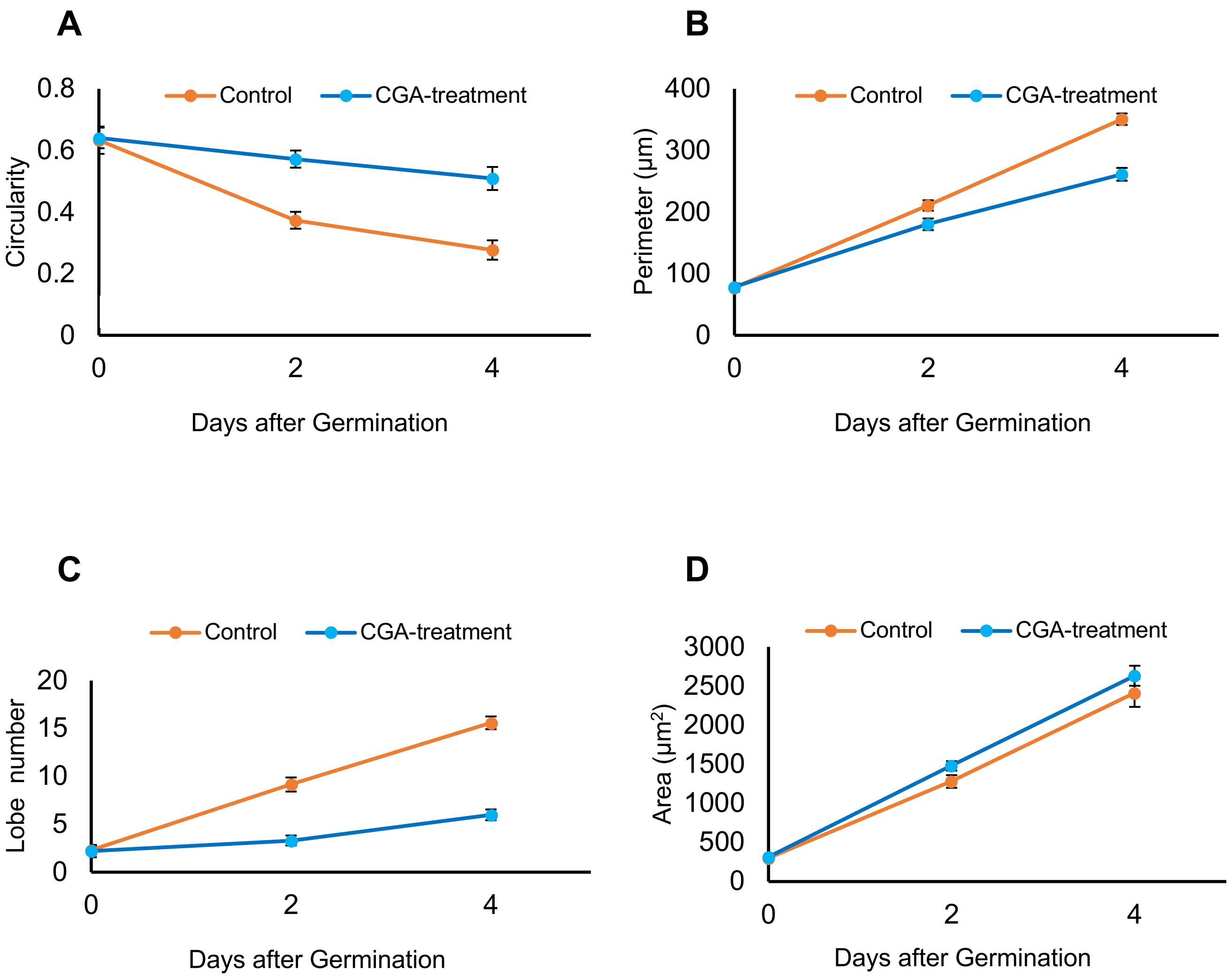


**Figure S8.** Related to Figs. 6H-K, **effect of CGA treatment on pavement cell growth and lobiness**. Comparison of the pavement cells on adaxial surfaces of cotyledons of *Arabidopsis thaliana* seedlings grown in presence of CGA to reduce the cellulose crystallinity (control samples contain corresponding amount of DMSO). Pavement cells in treated samples showed reduced ability to form wavy borders **A)** and **B)**. **C)** and **D)** show that both group of samples grow similarly in terms of area and perimeter expansion and the reduced number of lobes was not a consequence of growth arrest in treated samples. Error bars show standard error.
